# Facilitate integrated analysis of single cell multiomic data by binarizing gene expression values

**DOI:** 10.1101/2024.02.22.581665

**Authors:** Rohan Misra, Alexander Ferrena, Deyou Zheng

## Abstract

The identity of a cell type can be revealed by its transcriptome and epigenome profiles, both of which can be in flux temporally and spatially, leading to distinct cell states or subtypes. The popular and standard workflow for single cell RNA-seq (scRNA-seq) data analysis applies feature selection, dimensional reduction, and clustering on the gene expression values quantified by read counts, but alternative approaches using a simple classification of a gene to “on” and “off” (i.e., binarization of the gene expression) has been proposed for classifying cells and other downstream analyses. Here, we demonstrate that a direct concatenation of the binarized scRNA-seq data and the standard single cell ATAC-seq data is sufficient and effective for integrated clustering analysis, after applying term-frequency-inverse document frequency (TF-IDF) and single value decomposition (also called latent semantic indexing, LSI) algorithms to the combined data, when the two modalities of omic data are collected using paired multiomic technology. This proposed approach avoids the need for converting scATAC-seq data to gene activity scores for combined analysis and furthermore enables a direct investigation into the contribution of each data type to resolving cell type identity.

## Introduction

Single cell omic analysis has been adopted widely for resolving cellular heterogeneity existing in biological samples, leading to novel insights to cell differentiation, tissue/organ development, and cancer formation ^1,2,3^. A key step in the data analysis is cell clustering, based on between-cell similarity of the underlying quantitative data, for example, gene expression profiles. Not only is clustering important for distinguishing cell types (or states), but also critical for almost all the downstream bioinformatics analysis. Moreover, investigators have become increasingly interested in collecting and analyzing multiple types of omics data, either from the same cells or different cells of biologically matching samples. Among the technologies, single cell RNA-seq (scRNA-seq) is more advanced, followed by single cell ATAC-seq (scATAC-seq). Commercial platforms (e.g., from the 10X Genomics) are also available for capturing single cells for simultaneously interrogating the gene expression profile and chromatin accessibility of the same cells, yielding paired multiomic data.

Many software packages have been developed for analyzing these two modalities of data and their integration^2^. A common analytic workflow will analyze and cluster scRNA-seq data and scATAC-seq data separately and then either project the scRNA-seq clustering information to the scATAC-seq data (e.g., by label transfer), if the two data types are collected independently, or convert the scATAC-seq data to “gene activity scores” for combined analysis ^4,5^, if data from the two modalities are collected using a multiomics platform ^6, 7^. However, the method for projection of scATAC-seq data to the gene space is rather subjective, which basically sums scATAC-seq reads within a distance to a gene based on chromosomal coordinate, and thus should be improved if possible. Consequently, such a strategy only uses part of the scATAC-seq data, potentially leaving out critical information. Whether the two types of data can be combined directly for integrated analysis, to our knowledge, has not been attempted. A concern may be that scRNA-seq quantifies gene expression by RNA abundance, which can vary in several orders of magnitudes between highly expressed genes (up to hundreds of reads) and lowly expressed ones, while scATAC-seq quantifies chromatin accessible or not, with values in low single digits (95% of measurements < 2; **Figure S1**). Another concern may be related to the fact that scATAC-seq data is sparser than scRNA-seq data, even though both are very sparse with “0” entries for most cells.

The former concern can be addressed by transforming the scRNA-seq data to a binary format, by defining the expression of individual genes as on (thus “1”) and off (thus “0”) in individual cells. Previously, binarization of scRNA-seq data has been applied successfully to cell clustering ^8,9^, differential expression analysis ^10^, and other downstream analysis, such as inferring cell developmental trajectory ^8^, gene regulatory network ^11^, and cell age ^12^. In this study, considering the similarity between binarized scRNA-seq data and scATAC-seq data, we first addressed how the popular algorithms used in scATAC-seq analysis, specifically TF-IDF/LSI ^13^, could be applied to binarized scRNA-seq data for improving cell clustering. After that, we demonstrated that the combined matrix containing scATAC-seq data and binarized scRNA-seq data (from paired multiomic assays) could be analyzed and clustered using current scATAC-seq workflow, yielding clustering results highly similar to or better than what were obtained by standard integration methods, which need more complex algorithms and more computing resources. Additionally, we were able to study how the two data modalities affect the separation of specific cell types in some datasets by changing the RNA to ATAC features. In short, our study suggests that the binarized and concatenated approach holds promising values in integrated analysis of multiomic data and probably should be considered as a baseline method before using more complex algorithms.

## Data and Methods

### Processing and clustering of scRNA-seq data using standard workflow

Two popular data analysis platforms were explored, based on either SCANPY ^14^ (Python) or Seurat software (R) ^15^. We refer to the general scRNA-seq analysis workflow outlined in the Scanpy tutorial ^14^ and implemented in the Scanpy (v1.9.3; Python v3.10) as the “standard workflow.” Likewise, the general workflow in Seurat was considered the “standard workflow.” In both cases, we used default parameters unless specified. These standard workflows included quality control (QC) to filter out poor quality cells and genes detected in a few cells, read count normalization and scaling, highly variable feature (HVF) selection, and dimensionality reduction by principal component analysis (PCA). In the Scanpy workflow, we also regressed out the effect of mitochondrial genes by the “regress_out” function, while 30 PCs (unless specified otherwise) were used for dimensionality reduction, neighborhood detection, clustered by the Leiden algorithm (v0.9.1)^16^, and visualized by either Uniform Manifold Approximation and Projection (UMAP) or t-distributed stochastic neighbor embedding (t-SNE). Batch correction, if required, was achieved by utilizing the Batch Balanced KNN algorithm (v1.5.1)^17^. In the Seurat workflow, for non-binary data analysis, Seurat (v5.0.1) ^15^ was used with the SingleCellTransform ^18^ workflow and default parameters. Elbow plots were used to determine the optimal number of PCs. Graph construction, UMAP visualization, and Louvain clustering were used with data-specific PC and resolution parameters as described in more detail in Supplement.

### Processing and clustering of scATAC-seq data using standard workflow

We followed the standard scATAC workflow of MUON^19^ (v0.1.5; Python v3.10; part of scverse ecosystem ^20^ ) as outlined in its documentation for analyzing scATAC-seq data. For QC, ATAC peaks detected in less than 10 cells were removed with the “filter_vac” function, as were cells with less than 2,000 genes, by the “filter_obs” function. The resulted peaks by cells matrix was then normalized using term-frequency inverse-document-frequency (TF-IDF), followed by singular value decomposition (SVD) for dimensionality reduction in Muon. The TF-IDF/SVD is known as Latent Semantic Indexing (LSI), but some also use LSI for the dimensionality reduction step after TF-IDF, e.g., in Muon. The resulted quantitative matrix was then processed using the standard Scanpy workflow described above, including visualization in t-SNE or UMAP, and clustering by the Leiden algorithm.

### Processing and clustering of binarized scRNA-seq data using scATAC-seq workflow

Binarization of scRNA-seq data was accomplished by thresholding the gene expression matrix (after QC filtering described above), converting any value above zero to 1, while the rest as 0. The resulted binary matrix was then processed as described above for scATAC-seq data matrix, including normalized by TF-IDF and LSI in Muon. The subsequent clustering and low dimensional visualization also followed the scATAC-seq workflow.

Additionally, we applied the Seurat/Signac workflow on binarized scRNA-seq data and combined scRNA-seq and scATAC-seq data. The R package Signac ^21^ (v1.11.0) was used with default settings for LSI (TF-IDF/SVD) analysis. Elbow plots and “depth correlation” plots were used to inspect LSI components. LSI component 1 was always excluded. Graph construction, UMAP visualization, and Louvain clustering were used with data-specific PC and resolution parameters as described in Supplement.

### Datasets and data-set specific processing

Human pancreatic islet scRNA-seq data (accession numbers: E-MTAB-5061and E-MTAB-5060) were obtained from a previous study described in Segerstolpe et al. ^22^. It contained 2,209 cells from both healthy and type 2 diabetic donors and was generated by the Smart-seq2 technology. The expression data in either raw count or RPKM (Reads Per Kilobase of transcript per Million mapped reads) form were used, while the cell cluster and corresponding type annotation from the authors were used without modification in this study.

Two sets of data from human peripheral blood mononuclear cells (PBMC) were used. One is scRNA-seq data from 3,000 PBMCs (referred as “3K”), downloaded from the 10X Genomics^23^. This dataset has been used as a standard for many software development, including Seurat and Scanpy ^15,24^ . The other is a multiomic dataset provided by the 10X Genomics and contains scRNA-seq and scATAC-seq data from 11,909 PBMCs (referred as “10K”) ^25^. In the ATAC-seq portion, cells contained a median of 13,486 high quality ATAC peaks. Standard cell type clusters were not available for this dataset, so we used the previously identified canonical markers to assist our clustering and cell type annotation ^26^.

A large dataset comprising of 45,870 cells, 78,023 CD45^+^ enriched cells and 363,213 nuclei from 14 adult human hearts was obtained from a previous study ^27^ (Human Cell Atlas Accession no. ERP123138). It has a total of 487,106 cell/nuclei barcodes and 11 major cell types. We used the filtered and normalized data provided by the authors, as well as their cell clusters and annotations for identifying cells.

A dataset from the human cerebral cortex consisting of single-nuclei paired RNA- and ATAC-seq of 45,549 nuclei prepared via the 10X Multiome platform was also used ^28^ . Count matrices were downloaded from the GEO (Accession “GSE204684”) while cell annotation metadata was from the Broad Single Cell Portal under accession “SCP1859”. The analysis of this dataset used the workflow implemented in Seurat and Signac (see Supplement).

A dataset from non-hematopoietic bone tissue comprising the bone marrow stromal niche consisting of single-cell RNA-seq of 20,581 cells prepared via the 10X Chromium v2 platform was used ^29^. The expression count matrix and cell annotation metadata were obtained from the Broad Single Cell Portal under accession “SCP361”.

Finally, the mouse breast multiomic data were described in a previous publication by Foster et al ^30^. The study analyzed both human and mouse normal breast tissues and breast cancer samples using the 10X Multiomic platform, but we used only one of the samples from the normal non-tumor mammary parenchyma of mouse breast tissue (GEO: GSM6543819). After QC, we obtained 9,057 cells with 25,767 genes from scRNA-seq and 93,484 fragments/peaks from scATAC-seq for our integration study.

## Results

### Clustering single cells using binarized gene expression data

Before going directly to integrated analysis of scRNA-seq and scATAC-seq data, we wanted to evaluate the usage of binarized scRNA-seq data for cell clustering. We began by studying how the clusters from binarized data match to those from a standard Scanpy workflow ^24^ without applying TF-IDF (**Figure 1)**. We started with a 3K PBMC scRNA-seq dataset ^23^ that has been frequently used for benchmarking scRNA-seq software performance ^24,31^ . After standard quality control of the raw count data (i.e., removing cells of poor quality and genes expressing in few cells), we binarized the input scRNA-seq data with read counts (referred as “pre-binarized RNA” data) by converting the expression values to 1 if the raw read count was > 0, and otherwise 0 (see Methods). Using 2,000 top highly variable genes, 30 principal components (PCs) and otherwise default parameters in Scanpy, we obtained 8 clusters from the binarized data by Leiden clustering (resolution = 1.0) (**Figure S2A**,**B**). As the cell types were available for the data, we compared our clusters to the annotated cell types and found that > 77% of cells in our clusters (ranging from 44% to 100%) were from a single PBMC cell type, except for CD4+ T cells and CD14 monocytes, which were each split into two large clusters (**Figure S2C**). Two of the clusters (#1 and #4) both contained two cell types (CD14+ Monocytes and FCGR3A Monocytes; CD8+ and NK cells) but the mixed cell types were relatively harder to distinguish than other cell types, as indicated by the expression of the canonical gene markers (**Figure S2B**). Using the same method and parameters, we binarized and clustered a human pancreatic cell dataset obtained by the Smart-Seq2 platform^22^, yielding 14 clusters (**Figure S2D**,**E**). In this case, over 74% of cells (ranging from 44% to 100%) were clustered in a manner highly consistent with the authors’ original cell annotation, except for alpha and beta cells, which were split into 4 and 2 clusters, respectively (**Figure S2F**). Multiple very small clusters, however, mistakenly combined cells from distinct cell types. Overall, these results confirm previous reports that binarized scRNA-seq data could be directly used for cell clustering, but some methodology improvement is needed ^8, 9^.

**Figure 1.**
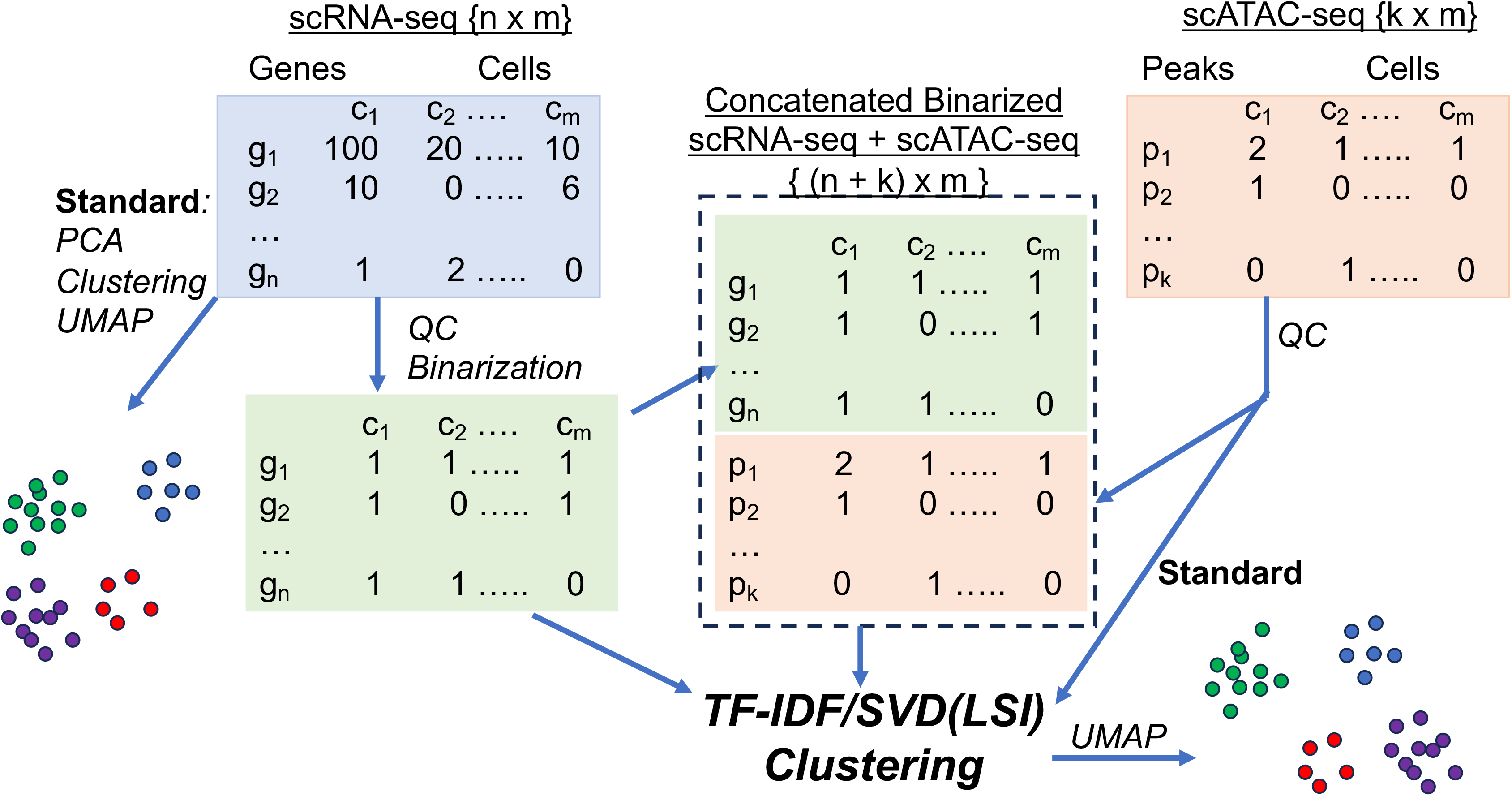
Flowchart delineating the steps using binarized scRNA-seq data for cell clustering and integrated analysis with scATAC-seq data.

Considering the high similarity between binary scRNA-seq data and scATAC-seq data matrix in terms of sparsity and small (count) values, we decided to test whether the standard scATAC-seq analysis algorithm can be applied, i.e., subject the binarized scRNA-seq matrix to normalization by term-frequency inverse-document-frequency (TF-IDF), dimensionality reduction by SVD, and clustering by Leiden algorithm using highly variable genes selected by analytical Pearson residuals (**Figure 1**). The TF-IDF-SVD is often called Latent Semantic Indexing (LSI)^32^, but sometime LSI refers to the dimensionality reduction step, thus we use TF-IDF/LSI. Interestingly, this indeed led to improved clustering of the 3K PBMC data, with the mean accuracy increased to 86% (**Figure S3A-C**). Moreover, the CD8+ T and NK cells were separated correctly. So were CD14+ Monocytes and FCGR3A Monocytes. For the human pancreas dataset, the binarization protocol was followed by an integration step using Batch Based KNN (K nearest neighbor) ^17^ to reduce the batch effects due to donors. After that, we obtained 14 clusters (**Figure S3D-F**), the same as reported by the original authors ^22^. For the pancreatic dataset, over 94% of total cells were clustered with their own cell type. We further evaluated our TF-IDF/LSI approach on a much larger dataset, containing 486k cell/nuclei barcodes from adult human hearts ^27^. Binarization and clustering yielded 25 clusters (**Figure S4A**), while the original study reported 13 (11 cell types + “doublets” and “unassigned”). Comparing the expression of the marker genes indicated that the split of multiple cardiac cell types in the binarization clusters reflect the difference in the percentages of cells expressing those markers, e.g., clusters 3, 4, and 5 were all ventricular cardiomyocytes but cluster 3 displayed a smaller % of cells expressing *MYH7* (**Figure S4B**). The myeloid and lymphoid cells were also each split into 2-3 clusters (**Figure S4C**). We further compared our clusters to the cell subtypes for ventricular cardiomyocytes (vCM) and endothelial cells (EC) ^27^, but did not find a simple concordance (**Figure S4D**,**E**).

To demonstrate that the clustering improvement from TF-IDF/LSI is not software dependent, we performed the binarization and clustering of a mouse bone cell atlas data using Seurat ^15^ and Signac ^21^ (both implemented in R). When compared to the authors’ original cell types, on average 88% of the cells in each of our cluster were from a single cell type (**Figure S5**).

Taken together, these results indicate that eliminating quantitative information related to gene expression magnitude does not remove signals pertinent to the distinct function of individual cell types, and that processing the binarized scRNA-seq data with the TF-IDF/LSI method used in scATAC workflow provides a marked improvement in clustering quality.

### Integrated analysis of binarized scRNA-seq data with scATAC-seq data

A key rationale for using TF-IDF/LSI to binarized scRNA-seq data lies in its potential for direct integrated analysis of scRNA-seq and scATAC-seq data that are acquired from the same cells using a multiomic platform. For this, the binarized scRNA-seq data matrix is directly concatenated with the scATAC-seq peaks x cells matrix, generating “BC” (for binarized and concatenated) matrix. This is feasible because both types of data are from the same cells and the columns of the two matrices (i.e., cell barcodes) are thus the same. The BC matrix is then processed as described above for scATAC-seq data (**Figure 1**). To test the performance, we obtained and analyzed a paired multi-omic10K PBMC data provided by 10X Genomics. We first performed clustering on the pre-binarized scRNA-seq data (**Figure 2A**) and the scATAC-seq data (**Figure 2B**), leading to 13 and 12 clusters, respectively. Analyses of the expression (or chromatin accessibility) patterns of the marker genes identified previously for the same data ^26^ allowed us to successfully assign the cell type to each cluster (**Figure S6A-B**). We also clustered the binarized scRNA-seq data (**Figure S7A)**. Clustering of the combined scRNA-seq and scATAC-seq data yielded 21 clusters (**Figure 2C**), with several cell types being split into multiple clusters as indicated by the expression of the cell type markers (**Figure S6C**). Comparison of the clusters from the scRNA-seq or scATAC-seq data with the integrated clusters (referred to as “BC” clusters) further showed that five of the 13 cell types were split to more than one clusters in the integration result (**Figure 2D,E**), as expected from increased clustering resolution. Interestingly, 4 BC clusters (0, 6, 7, 13) all contained a mixture of two cell types, CD14 monocytes and intermediate monocytes, when compared to the clusters based on scRNA, but this problem did not occur when compared to the clusters generated with scATAC data only. Thus, we compared the clusters from the two modalities before integration and found a large disagreement in their clustering of these two cell types (**Figure S7B-C**). Nevertheless, >80% of the cells in the majority of the 21 BC clusters were from a single cell type defined by either scRNA or scATAC data modal (**Figure 2F**; **Figure S7D**,**E**).

**Figure 2.**
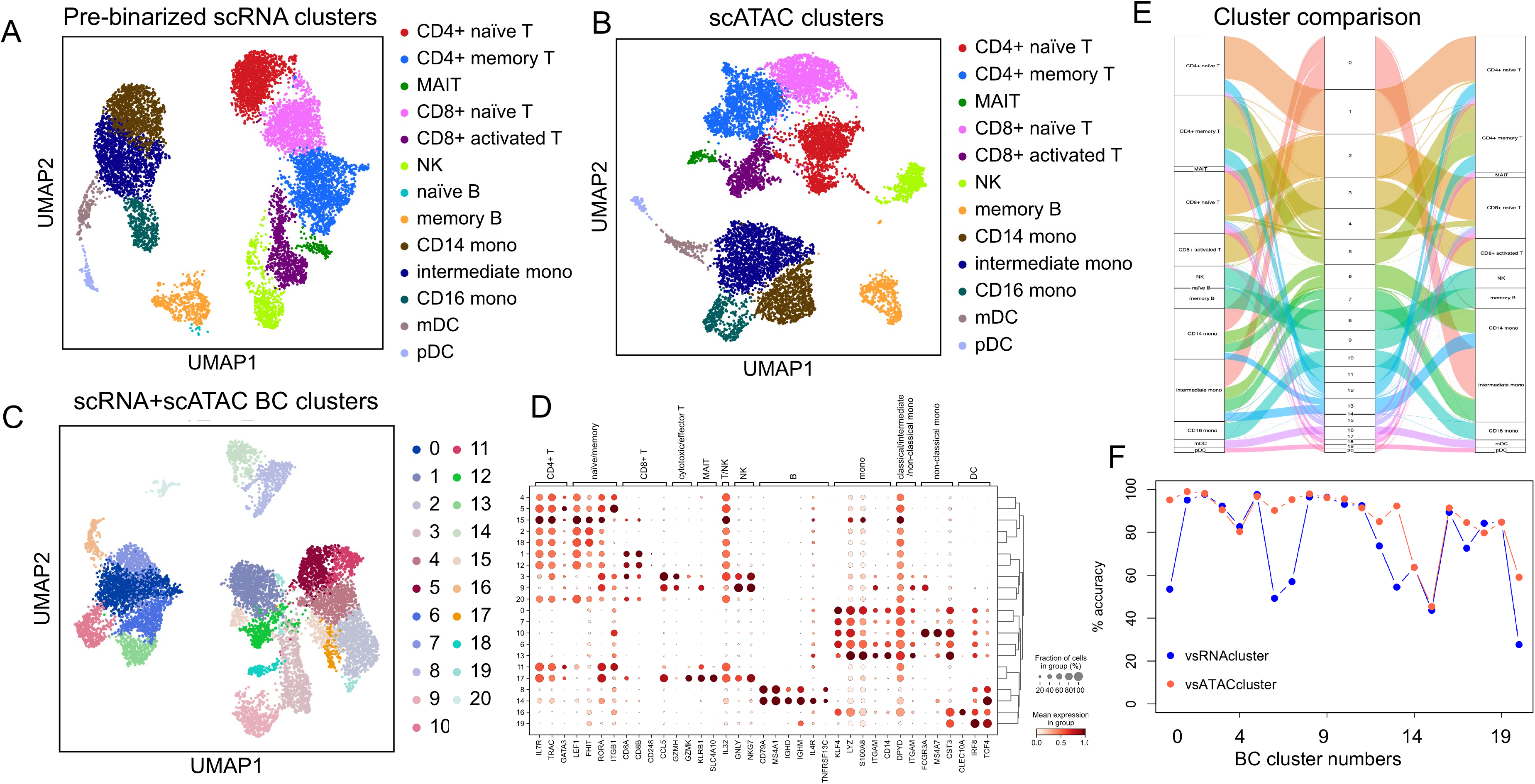
Clustering concatenated scRNA-seq binarized data and scATAC-seq 10K PBMC data. **A**). UMAP visualizing of the pre-binarized scRNA-seq data, colored by Leiden clusters with our re-annotated cell types. **B**). UMAP of the scATAC-seq data, colored by re-annotated cell types. **C**) Integrated clustering of the binary scRNA-seq data and the scATAC-seq data, colored by clusters. The number of BC clusters were chosen to match that from MOFA integration (Figure 3). **D**). Bubble plot showing the expression of the cell type markers previously used to annotate cell types. Note that the expression in all bubble plots was from pre-binarized data. **E**). Alluvia plot showing the cell relationship between cell types in A (left) the clusters in C (middle), and the clusters in B (right). **F**). Cluster accuracy, as determined by the %s of cells in the dominant cell type determined in (A) or (B) for each of the integrated BC clusters (x-axis).

We performed a cluster marker identification from the BC matrix using Scanpy (“rank_genes_groups” function; adjusted p < 0.05 and log2(fold change) > 0.5; t-test) and found that the ratios of genes to ATAC peaks among the markers varied significantly across the 21 clusters, from 0.03 to 0.31 (excluding the outlier “CD8_Naive_2”) whereas the ratio for the total features in the BC matrix was 0.25 (**Table S1**). Moreover, a significant fractions of marker peaks were not linked to the marker genes in the same clusters (**Table S1**), supporting the value of our integration analysis.

After showing that binarized scRNA-seq data and scATAC-seq data could be directly combined for clustering analysis, we set out to compare the clustering results from our integrated approach to those using the non-binarized multi-omic data. We applied the MOFA algorithm ^33^ to integrate the 10K PMBC multiomic data, followed by nearest-neighbor graph construction, UMAP calculation and Leiden cluster. The result showed that clustering results from the two methods are mostly comparable, when the cells were clustered into the same number of groups (**Figure 3A-C**). A key difference is that some intermediate monocytes in the BC cluster 0 were split into 5 clusters by MOFA (**Figure 3D**,**E; Figure S7F**), but both approaches split intermediate monocytes to multiple clusters (**Figure 2,3**), suggesting intermediate monocytes have high transcriptomic diversity. Such a difference in cluster splitting was observed for clusters 6, 7 and 13 too. When the qualities of the Leiden-based clusters were assessed by the Silhouette scores ^34^, clusters from both methods had comparable scores (0.33 vs 0.38), indicating similar compactness. Using the cell types defined by clustering the pre-binarized scRNA-seq data (**Figure 2A**), we also computed the Adjusted Rand Index (ARI) ^35^ for the clusters and obtained comparable values (0.53 for BC vs 0.58 for MOFA). Since the ARI is a quantification for how often cells of the same type are correctly clustered, and both integration methods yielded very similar results, we concluded that integration using BC data is a valid approach. We should note that cluster numbers would affect ARI and we had significantly more clusters than the 13 cell types, because for the purpose of comparable we have set the parameters in clustering our BC data (Leiden resolution = 1.0) to match the cluster number from MOFA integration using their default parameters (Leiden resolution = 0.9).

**Figure 3.**
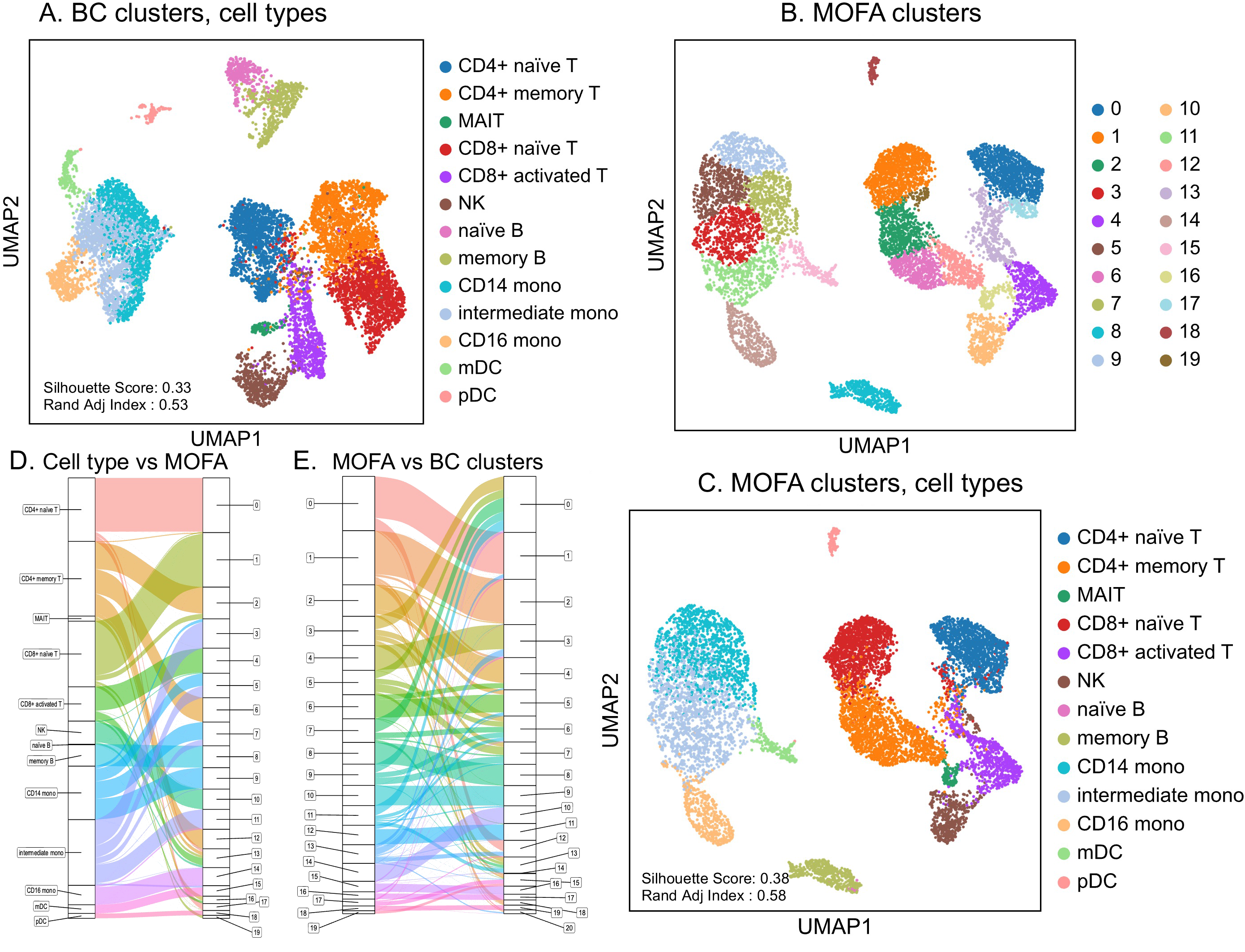
Comparison of the clustering results from MOFA integration and from our BC approach. **A**) UMAP of BC clustering results as in Figure 2C but here colored with known cell types. **B-C**) UMAPs of the clustering result of the MOFA2+ integration, colored by Leiden clusters (B) or cell types (C). **D-E**) Alluvial plots showing the relationship of known cell types with the MOFA integration clusters (D) and the relationship between MOFA and BC clusters (E).

### Altering the ratio of RNA and ATAC features for integration analysis

An important reason for collecting multiomic data is that the two data modalities provide complementary information and thus better for resolving cell types, especially subtypes. We examined the top 25,000 highly variable features (HVFs) from the combined BC data matrix, which contained features from both RNA and ATAC modalities (**Figure 4)**. Interestingly, the ratio of the RNA to ATAC features changed as more HVFs were selected, with RNA more prevalent among the top features (**Figure 4A**), suggesting that RNA may be more informative, despite the total number of ATAC features (i.e., peaks) being much more than the RNA features (i.e., genes) in this PBMC data.

**Figure 4.**
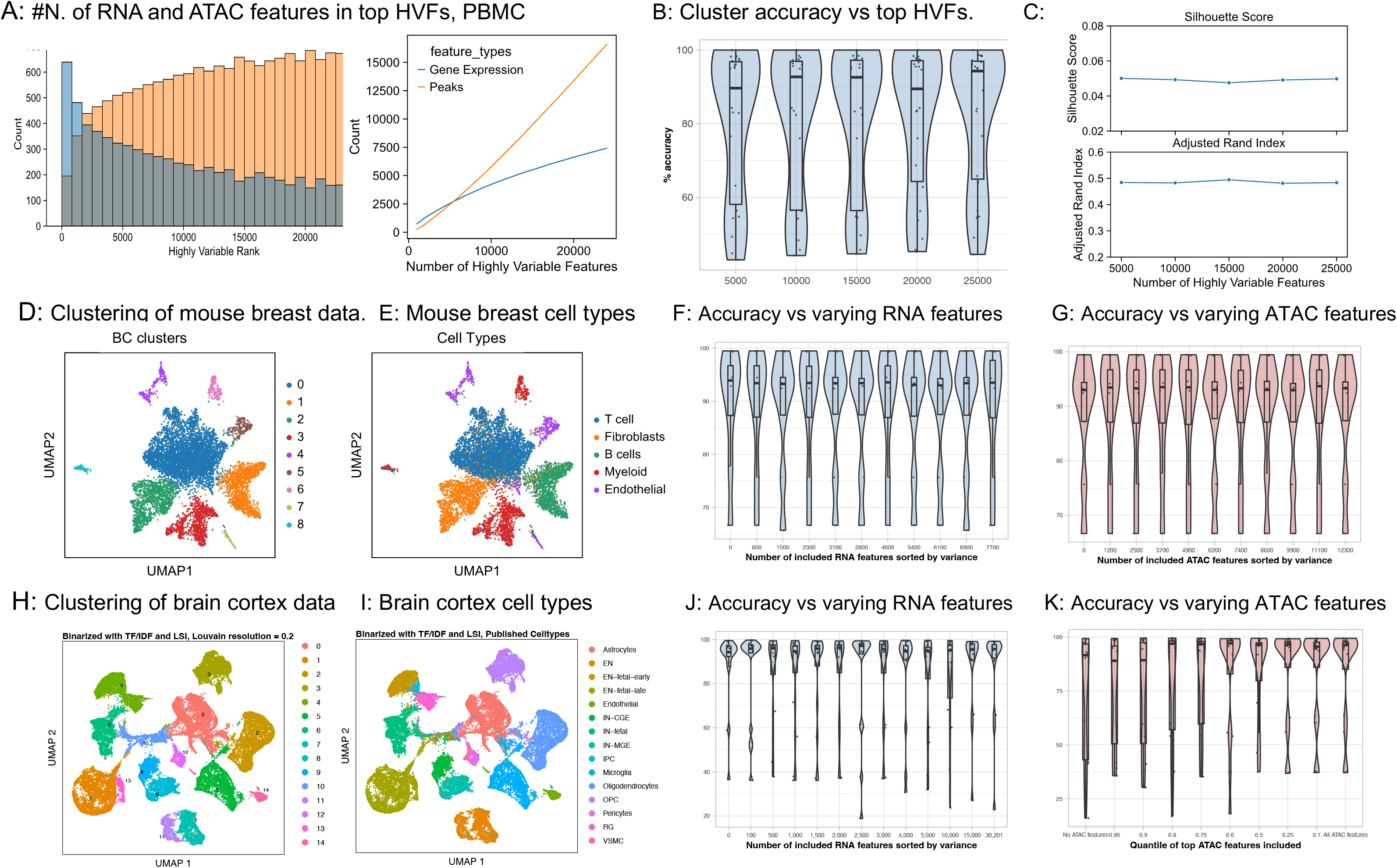
Effects of varying RNA:ATAC features on BC cell clustering. **A**) Bar plot shows the top 25,000 HVF in the binary integrated PBMC dataset, separated to features from RNA or ATAC modalities. **B**). Clustering accuracy with respect to the number of top HVFs. **C**). Silhouette scores and ARIs for clusters using different numbers of HVFs. **D,E**). UMAPs showing the 10X Multiome data from a normal mouse breast sample analyzed by our BC approach, with cells colored by BC clusters (D) or cell types (E). **F,G**). Violin plots showing the clustering accuracies when different numbers of RNA features (F) or ATAC features (G) were included in clustering the mouse breast cells. (**H, I**) UMAPs showing a human cortex 10X Multiome data analyzed by our BC approach, with cells colored by Louvain clusters (H) or the published cell types (I). **J,K**). Violin plots showing the clustering accuracies when different numbers of RNA features (J) or ATAC features (K) were included in clustering the human cortex cells. The middle lines in boxplots indicate median and the box lines represent 25^th^ and 75^th^ percentile of the data.

This observation prompted us to study how the ratio of the two features could impact the clustering results and moreover if some cell types were more affected. This was further motivated by our findings that (i) scRNA and scATAC data seemed to provide conflicting clustering for CD14+ monocytes and intermediate monocytes (**Figure S7C**), (ii) a main difference between BC clusters and MOFA clusters is that four of our BC clusters (0, 6, 7, 9) were split in the latter, such that our clustering result matched clusters based only on ATAC better while MOFA result matched clusters derived from only RNA better, and (iii) the ratios of marker genes to marker peaks varied significantly among cell types (**Table S1**). We decided to use top 5,000 to 25,000 HVFs in 5,000 increment to cluster the PBMC data. The results indicated that the ratios of RNA to ATAC features had different impacts on the cluster accuracies, with the clusters containing monocytes cells being affected more than others (**Figure 4B**). However, globally the distinctions from using different numbers of RNA and ATAC HVFs were overall not obvious, supported by Silhouette scores and ARIs (**Figure 4C**).

To explore further, we applied our BC clustering approach to a second multiomic data from mouse breast tissues (mammary parenchyma) ^30^, generating 9 clusters (**Figure 4D**). The authors identified five major cell types in the sample but did not share publicly the annotation for individual cells. Using the information as guidance, however, we were able to assign cell types to the 9 clusters based on canonical markers (**Figure 4E**). We then fixed the number of top ATAC HVFs (n = 12,300 from our full analysis) and added in gradually increased numbers of RNA HVFs for BC clustering. While increased usage of RNA features did not impact the clustering performance dramatically, the accuracies, however, seemed to increase and then decrease slightly (**Figure 4F; Figure S8**). Conversely, we fixed the top RNA HVF number (n = 7,700) and added in different numbers of top ATAC HVFs for clustering. As shown in **Figure 4G and Figure S8**, we found a gradual decrease in the mean clustering accuracies, suggesting for this dataset the ATAC data may be less informative. Interestingly, we found that some cell types were more affected than others by the RNA:ATAC ratio (e.g., endothelial cells and myeloid cells) (**Figure S9**). Moreover, we were able to assign the Endothelial-1 cluster as epithelial cells, as the cluster was *Epcam* and *Muc1* positive, while the Myeloid-3 and Myeloid-1 clusters most likely contained dendritic cells and macrophage (*CD68*+), when more features were used (**Figure S8**).

Finally, we tested the effects of varying top RNA to ATAC ratio using the Seurat/Signac workflow on a human cerebral cortex data ^28^, in this case using the authors’ weighted nearest neighbor integration of the RNA and ATAC as the reference clusters since they were available. The results support that the BC clustering accuracies could be affected by RNA to ATAC ratios (**Figure 4H-J**). Close examination of the results found that some brain cell types were affected significantly more than others (**Figure S10**); again, some cell types (or clusters) became more separated with more RNA or ATAC features included (e.g., inhibitory neuron subtypes) (**Figure S11**).

Taken together, our results indicate that the scRNA-seq and scATAC-seq BC data provides a good means for investigators to adjust the contribution of the two data modalities in their analysis, which in our view has the advantage in making better usage of the multiomic data and addressing which of the two types of omic data plays more important roles in defining the cell types of their interest.

## Discussion

Cell clustering is a fundamental step in single cell omic data analysis. In this study, consistent with previously reports, we found that applying TF-IDF/LSI algorithms to binarized scRNA-seq data could obtain clustering results comparable to what were obtained with non-binary data. While this may sound surprising, it has been reported before and the finding may be explained by this possibility: cells are clustered by similarities in their gene expression profiles, but given the sparsity of the data the similarities may have more to do with gene co-detection (i.e., co-expression) than the degree of their quantitative expression correlation, as suggested before ^8, 9^. Although more studies and further improvement are needed, our results also suggest that “dropout” (or zero counts), a commonly cited issue in single cell data, may not be as critical as discussed before ^36^, at least with respect to cell clustering. We should also note that TF-IDF transformation was previously tested on scRNA-seq data, in combination with various clustering algorithms, but the binarization step was different ^13^. From biology perspective, our finding suggests that a cell type is likely defined by a set of critical marker genes, whose activation is sufficient for its specification. The expression levels, on the other hand, may be important to define functional (or transcriptomic) states. More studies will be needed in the future to carefully address if these hold true. Additionally, it will be valuable to evaluate if other algorithms could perform better than SVD on the TF-IDF transformed BC matrix ^37^.

While performing analysis on a binary data matrix has advantages regarding computational efficiency, as described recently ^36^, the key advantage in our view, however, lies in the integration analysis of multiple types of omic data. In the current study, we only address scRNA-seq and scATAC-seq data from multiomic platform. The alternative and standard method is to compute gene activity scores from the scATAC-seq data and use them for integration. To do that, one needs to map individual scATAC-seq peaks to genes, often based on their distances on chromosomes. A caveat to consider here is that there are some peaks that may be associated to the wrong genes, as they may correspond to regulatory elements controlling genes far away from their locations. Furthermore, this method applied a cutoff to the physical distance, excluding many ATAC-seq data in the clustering. Our approach does not have this prediction step, as all genes and all peaks are considered together. Although we did not directly perform the analysis, it is conceivable that genes and ATAC peaks that cluster together are more likely to have a regulatory relationship than those that do not. Interestingly, for the 10K PMBC data, we found that the ratios of marker peaks associated with a marker gene of the same cluster to those without varied significantly among cell types and BC clusters, from 1.3 to 10.5 with a mean of 3.9 (**Table S1**). Moreover, a high percentage (15% on average) of marker peaks were more than 50 kb away from the assigned genes, regardless the genes were markers or not.

Using concatenation, data of other modalities can all be directly merged for integration analysis, after binarization if necessary, if all are acquired from the same cells. Note that we did not binarize scATAC-seq data, but the LSI was initially applied to the binarized scATAC-seq data^32^. Therefore, we believe binarization of scRNA-seq provides a logical and simple strategy for integrated analysis of multiomic data. Given its simplicity and interpretability, we would even recommend the BC approach to be considered as the baseline integration method before more complex algorithms are used. More importantly, as the scRNA-seq and scATAC-seq data are subject to the same computing process (TF-IDF/LSI, clustering, and visualization), it provides a good opportunity to investigate how these two modalities of data contribute to cell type identification. Although among the three datasets we tested, we did not see a significant change in overall cluster accuracy when more RNA or ATAC features were included. One possible explanation is that our starting point using only the top RNA or ATAC features already reached a high accuracy, and thus there was not much room for improvement. The other caveat of our results may be due to the lack of known clusters to be used as “gold standard” as all our performance metrics could be affected by the cluster numbers. Nevertheless, we did observe that the two types of data modalities affected cluster accuracies in a cell type dependent manner (**Figure S7-S11**). It will be valuable to test this on more datasets in the future.

Our binarization method leads to some information loss, and thus has limitations. Differential expression analysis is as important as clustering in single cell analysis. It is involved in marker gene identification and gene expression comparison across samples or conditions. The binary scRNA-seq data probably would not be suitable for these analyses, but they may be used to provide helpful information, for example, the fraction of cells in which a gene is expressed. The other limitation is that small cell clusters or cells with very few detected genes seem more likely to be split and merged into other cell clusters in the clustering of binary scRNA-seq data or after they were concatenated with scATAC-seq data. It would be important to investigate how this can be overcome in the future, or just to flag the cluster result for those cells as unreliable. Overall, we believe that our proposed binarization strategy has promising applications but needs other investigators’ inputs for further improvement.

## Data and code availability

All the scRNA-seq and scATAC-seq data were obtained from previous studies. The R and Python codes for data analysis will be made publicly available at GitHub.

## References

1. Stegle, O., Teichmann, S. A. & Marioni, J. C. Computational and analytical challenges in single-cell transcriptomics. Nat. Rev. Genet. 16, 133–145 (2015).

2. Heumos, L. et al. Best practices for single-cell analysis across modalities. Nat. Rev. Genet. 24, 550–572 (2023).

3. Stuart, T. & Satija, R. Integrative single-cell analysis. Nat. Rev. Genet. 20, 257–272 (2019).

4. Cusanovich, D. A. et al. A Single-Cell Atlas of In Vivo Mammalian Chromatin Accessibility. Cell 174, 1309-1324.e18 (2018).

5. Granja, J. M. et al. ArchR is a scalable software package for integrative single-cell chromatin accessibility analysis. Nat. Genet. 53, 403–411 (2021).

6. Miao, Z., Humphreys, B. D., McMahon, A. P. & Kim, J. Multi-omics integration in the age of million single-cell data. Nat. Rev. Nephrol. 17, 710–724 (2021).

7. Nam, A. S., Chaligne, R. & Landau, D. A. Integrating genetic and non-genetic determinants of cancer evolution by single-cell multi-omics. Nat. Rev. Genet. 22, 3–18 (2021).

8. Li, R. & Quon, G. scBFA: modeling detection patterns to mitigate technical noise in large-scale single-cell genomics data. Genome Biol. 20, 193 (2019).

9. Qiu, P. Embracing the dropouts in single-cell RNA-seq analysis. Nat. Commun. 11, 1169 (2020).

10. Bouland, G. A., Mahfouz, A. & Reinders, M. J. T. Differential analysis of binarized single-cell RNA sequencing data captures biological variation. NAR Genomics Bioinforma. 3, lqab118 (2021).

11. Moignard, V. et al. Decoding the regulatory network of early blood development from single-cell gene expression measurements. Nat. Biotechnol. 33, 269–276 (2015).

12. Yu, D. et al. CellBiAge: Improved single-cell age classification using data binarization. Cell Rep. 42, 113500 (2023).

13. Moussa, M. & Măndoiu, I. I. Single cell RNA-seq data clustering using TF-IDF based methods. BMC Genomics 19, 569 (2018).

14. Tutorials — Scanpy 0.1.0.dev documentation. https://scanpy.readthedocs.io/en/stable/tutorials.html.

15. Hao, Y. et al. Dictionary learning for integrative, multimodal and scalable single-cell analysis. Nat. Biotechnol. 1–12 (2023) doi:10.1038/s41587-023-01767-y.

16. Traag, V. A., Waltman, L. & van Eck, N. J. From Louvain to Leiden: guaranteeing well-connected communities. Sci. Rep. 9, 5233 (2019).

17. Polański, K. et al. BBKNN: fast batch alignment of single cell transcriptomes. Bioinformatics 36, 964–965 (2020).

18. Hafemeister, C. & Satija, R. Normalization and variance stabilization of single-cell RNA-seq data using regularized negative binomial regression. Genome Biol. 20, 296 (2019).

19. Bredikhin, D., Kats, I. & Stegle, O. MUON: multimodal omics analysis framework. Genome Biol. 23, 42 (2022).

20. Virshup, I. et al. The scverse project provides a computational ecosystem for single-cell omics data analysis. Nat. Biotechnol. 41, 604–606 (2023).

21. Stuart, T., Srivastava, A., Madad, S., Lareau, C. A. & Satija, R. Single-cell chromatin state analysis with Signac. Nat. Methods 18, 1333–1341 (2021).

22. Segerstolpe, Å. et al. Single-Cell Transcriptome Profiling of Human Pancreatic Islets in Health and Type 2 Diabetes. Cell Metab. 24, 593–607 (2016).

23. pbmc3k -Datasets -Single Cell Gene Expression -Official 10x Genomics Support. https://support.10xgenomics.com/single-cell-gene-expression/datasets/1.1.0/pbmc3k.

24. Wolf, F. A., Angerer, P. & Theis, F. J. SCANPY⍰: large-scale single-cell gene expression data analysis. Genome Biol. 19, 15 (2018).

25. pbmc_granulocyte_sorted_10k -Datasets -Single Cell Multiome ATAC + Gene Exp. - Official 10x Genomics Support. https://support.10xgenomics.com/single-cell-multiome-atac-gex/datasets/1.0.0/pbmc_granulocyte_sorted_10k?

26. Processing chromatin accessibility of 10k PBMCs — muon-tutorials documentation. https://muon-tutorials.readthedocs.io/en/latest/single-cell-rna-atac/pbmc10k/2-Chromatin-Accessibility-Processing.html.

27. Litviňuková, M. et al. Cells of the adult human heart. Nature 588, 466–472 (2020).

28. Zhu, K. et al. Multi-omic profiling of the developing human cerebral cortex at the single-cell level. Sci. Adv. 9, eadg3754.

29. Baryawno, N. et al. A Cellular Taxonomy of the Bone Marrow Stroma in Homeostasis and Leukemia. Cell 177, 1915-1932.e16 (2019).

30. Foster, D. S. et al. Multiomic analysis reveals conservation of cancer-associated fibroblast phenotypes across species and tissue of origin. Cancer Cell 40, 1392-1406.e7 (2022).

31. Hao, Y. et al. Integrated analysis of multimodal single-cell data. Cell 184, 3573-3587.e29 (2021).

32. Cusanovich, D. A. et al. Multiplex single-cell profiling of chromatin accessibility by combinatorial cellular indexing. Science 348, 910–914 (2015).

33. Argelaguet, R. et al. Multi-Omics Factor Analysis-a framework for unsupervised integration of multi-omics data sets. Mol. Syst. Biol. 14, e8124 (2018).

34. Rousseeuw, P. J. Silhouettes: A graphical aid to the interpretation and validation of cluster analysis. J. Comput. Appl. Math. 20, 53–65 (1987).

35. Hubert, L. & Arabie, P. Comparing partitions. J. Classif. 2, 193–218 (1985).

36. Bouland, G. A., Mahfouz, A. & Reinders, M. J. T. Consequences and opportunities arising due to sparser single-cell RNA-seq datasets. Genome Biol. 24, 86 (2023).

37. Zandigohar, M. & Dai, Y. Information retrieval in single cell chromatin analysis using TF-IDF transformation methods. in 2022 IEEE International Conference on Bioinformatics and Biomedicine (BIBM) 877–882 (2022). doi:10.1109/BIBM55620.2022.9994949.

