## Supplemental Figures for "Facilitate integrated analysis of single cell multiomic data by binarizing gene expression values"

Supplemental Results with Figures S1-11 Embedded.

### 1. Read counts for features in scATAC-seq data

We analyzed the values in the scATAC-seq data matrix of the 10K PBMC multiomics data and found that the majority of values (98%) were 2 or less (**Figure S1**), thus in the same scale of binarized scRNA-seq data.

**Figure S1. Accumulated distribution of fragment counts in scATAC-seq data.**

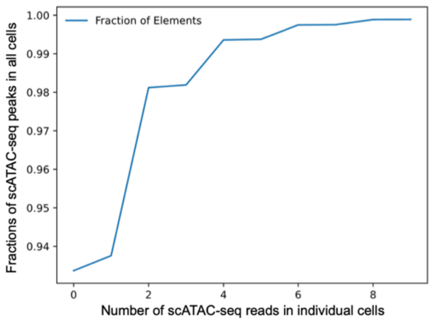


### 2. Binarizing and clustering human 3K PBMC and pancreatic scRNA-seq data using standard workflow

We binarized the scRNA-seq data for a human PBMC data generated by the 10X Genomics. It contained 2,700 and 2,698 cells before and after filtering, respectively. QC was done using default Scanpy parameters. The effects of total counts and potentially other confounders were removed by using the “regress_out” function of Scanpy against the calculated QC metrics of 'total_counts', 'pct_counts_mt', 'pct_counts_ribo', prior to binarization. The cell type annotation was obtained from the Muon documentation [1]. Highly variable genes (HVGs) were calculated on the binarized matrix using Scanpy’s highly_variable_genes function. Using the first 30 principal components, the neighborhood graph was constructed using the ‘neighbors’ function, and the cells clustered using the Leiden algorithm with a default resolution of 1.0. The clustering results of the binarized data are shown in **Figure S2A-C**. Similarly, we binarized and clustered a human pancreatic scRNA-seq data with cell type annotations provided by the authors (**Figure S2D-F**).

**Figure S2. Clustering scRNA-seq data after binarization.**

**A)** UMAP of the binarized 3K PBMC data, colored by clusters. **B)** Bubble plot showing the pre-binarized expression of canonical cell type markers for PBMC clusters in A. **C)** Matrix plot showing the cell relationship between clusters of binarized data and the cell types based on non-binarized PBMC data. **D)** UMAP of the binarized pancreatic data, colored by clusters. **E)** Bubble plot showing the pre-binarized expression of canonical cell type markers for pancreatic clusters in D. **F)** Matrix plot showing the cell relationship between clusters of binarized data and the cell types identified by the original authors.

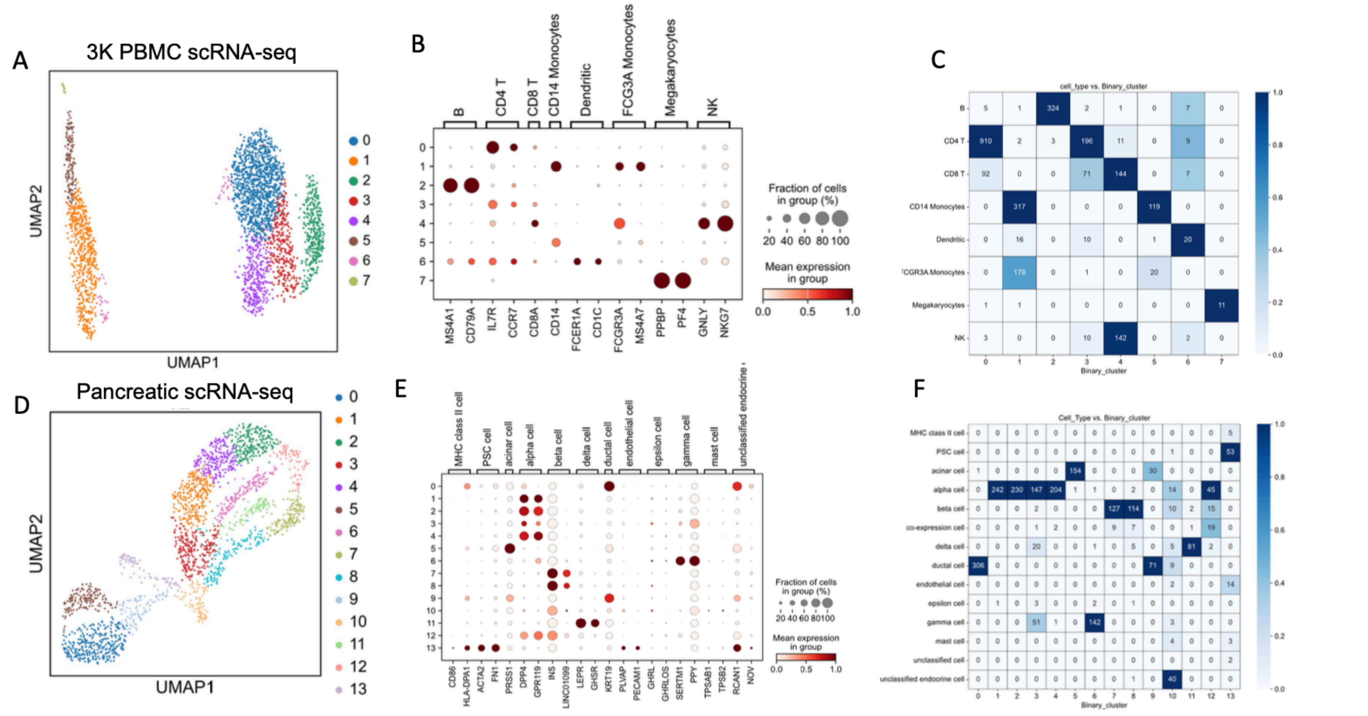


#### 3. Binarizing and clustering human 3K PBMC and pancreatic scRNA-seq data with TF-IDF transformation and LSI dimension reduction

Differently from the above, the binarized data were transformed by TF-IDF and subject to dimension reduction by LSI before clustering. The binarized matrix was transformed using the ‘tfidf’ function of muon, scaling to 10,000 counts per cell. Scanpy’s ‘normalize_per_cell’ function was subsequently used to normalize total counts per cell to 10,000. The matrix was then log transformed and scaled to 10. The LSI of the scaled matrix was calculated using Muon’s ‘lsi’ function. The first LSI component corresponded to the number of total read counts and thus was dropped. The next 30 LSI components were used for Neighborhood Graph construction and Leiden clustering (resolution = 0.9). For the pancreatic data, batch correction was performed using bbknn [2] . The results are in **Figure S3**.

**Figure S3. Clustering scRNA-seq data after binarization and TF/IDF transformation.**
**A)** UMAP of the binarized and TF/IDF 3K PBMC data, colored by clusters. **B)** Bubble plot showing the pre-binarized expression of canonical cell type markers for PBMC clusters in A. **C)** Matrix plot showing the cell relationship between clusters of binarized data and the cell types based on non-binarized PBMC data. **D)** UMAP of the binarized pancreatic data, colored by clusters. **E)** Bubble plot showing the pre-binarized expression of canonical cell type markers for pancreatic clusters in D. **F)** Matrix plot showing the cell relationship between clusters of binarized data and the cell types identified by the original authors.

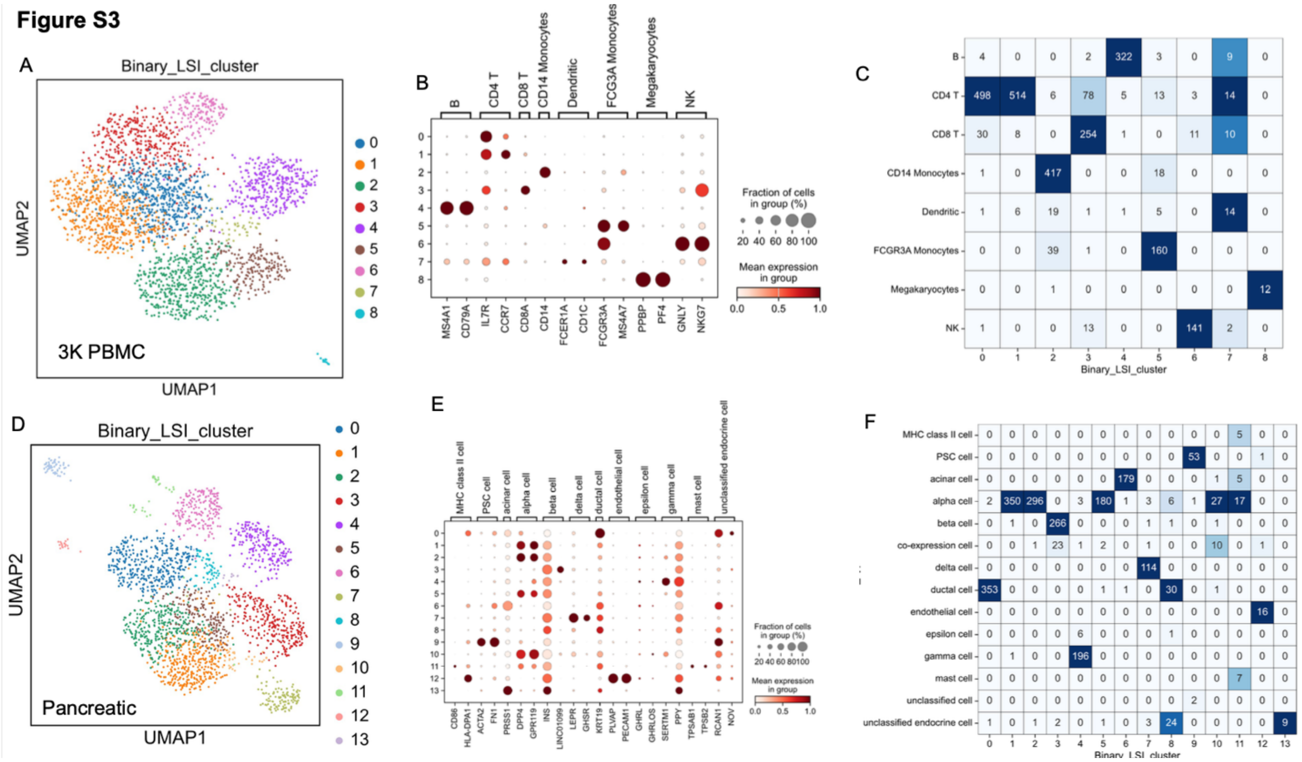

**4. BC clustering adult human heart scRNA-seq data with TF/IDF transformation and LSI dimension reduction**

We took a single cell (or nuclei) RNA-seq data from 14 human adult hearts that have been batched corrected. We binarized the expression matrix, applied TF/IDF transformation, LSI, and then Leiden clustering on the 2:31 dimension, as described above.

**Figure S4. Clustering heart scRNA-seq and snRNA-seq data.**

**A)** UMAP of the binarized and TF/IDF heart data, colored by clusters. **B)** Bubble plot showing the pre-binarized expression of cell type markers used by the original authors to identify main cardiac cell types. **C)** Matrix plot showing the cell relationship between our clusters of the binarized data and the main cardiac cell types defined by the original authors. **D)** Matrix plot showing the cell relationship between ventricular cardiomyocytes clusters of binarized data and the subtypes identified by the original authors. **E**) Matrix plot showing the cell relationship between endothelial clusters of binarized data and the subtypes identified by the original authors.

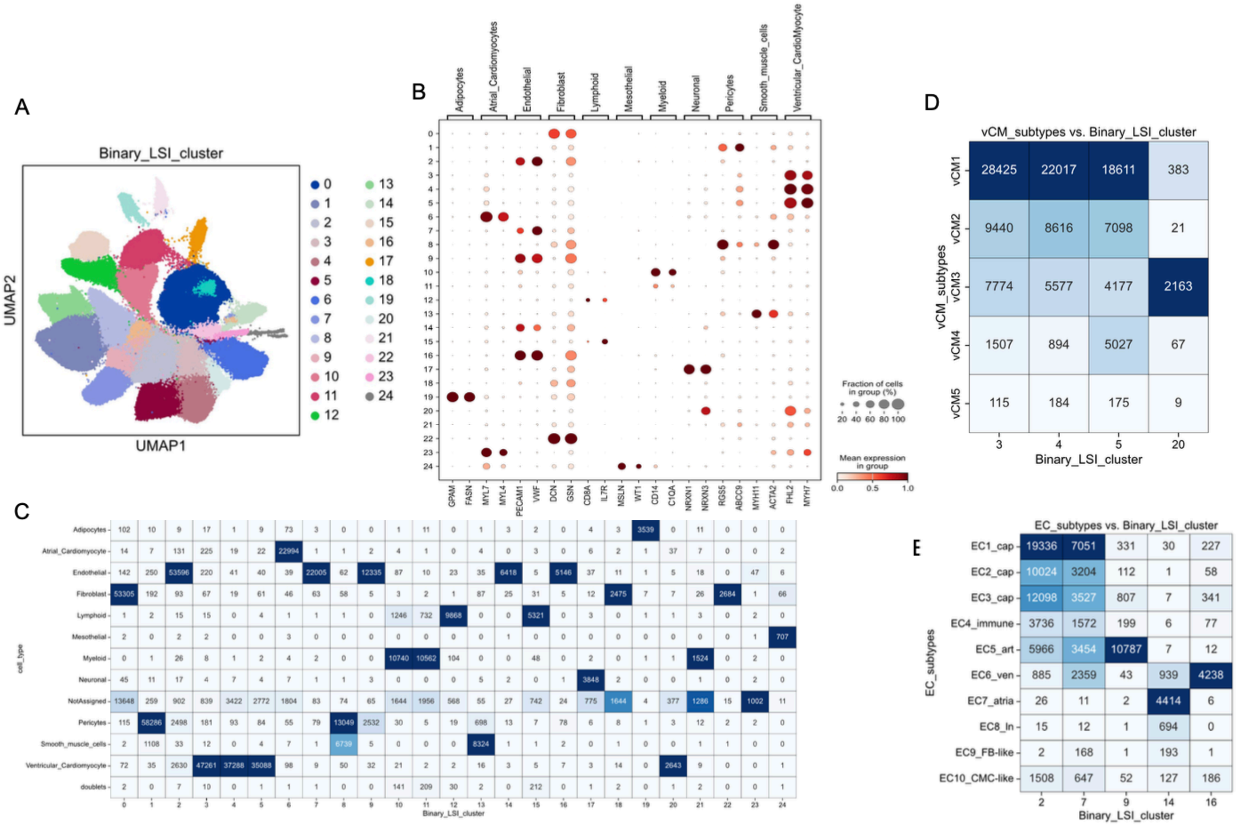

**5. Binarizing and clustering mouse bone atlas scRNA-seq data using Seurat software**

We tested our binarization approach on a mouse bone atlas scRNA-seq dataset, which contained 17 “cell types” by the authors’ annotation [3]. The data was downloaded from the Broad Single Cell Portal [4] (accession code “SCP361”). First, we performed Louvain clustering on the non-binarized expression data using Seurat [5] (v5.0.1) via the SCTransform workflows [5,6] (hvg =3000, npc =1:20, resolution = 0.5) and obtained 15 clusters (**Figure S5A**). The clusters were supported by the expression pattern of the top cell type markers identified by the authors previously (**Figure S5B**). We further used the authors’ cell type assignment to annotate our clusters, by dominance rule. The result showed high cluster concordance (**Figure S5C**) but with noticeable differences because some of the original cell types represented distinct cell states or subtypes (e.g., Fibro-4 and Fibro-5). Next, we binarized the expression matrix and performed Louvain clustering (hvg = 2000, npc = 1:15, resolution = 0.5), resulting in 12 clusters (**Figure S5D**). The expression of the cell type (or subtype) marker genes is shown in panel E (**Figure S5E**), indicating that different cell types were separated largely correctly but some subtypes were further merged or split, in comparison to our clusters from the non-binarized data (**Figure S5F**). Finally, we applied TF/IDF followed by LSI to the binarized matrix and used Louvain clustering (hvg=2000, npc = 2:10, resolution = 0.5), resulting in 11 clusters (**Figure S5G**). The expression of marker genes in these clusters are shown in **Figure S5H**, indicating that the clusters mostly recapitulated the cell types correctly.

To better define the cluster comparison, we analyzed the cell membership relation in the three approaches. For each set of our clusters, we first determined the dominant cell types, based on either the original author defined cell types (for the non-binarized data) or the clusters of the non-binarized data (for the binarized data), and then computed the %s of cells in the dominant cell type. By this metric, we found that the mean accuracy in our re-clustering of the non-binarized dataset was 84.2% (ranging from 43.1% to 100%) (**Figure S5C**), with the difference likely due to software choice and parameters. The mean accuracy in the clusters from the Louvain clustering of the binarized data is 87.1% (ranging from 64.1% to 98.2%) (**Figure S5F**), while the mean accuracy of the TF-IDF/LSI method is 88.0% (ranging from 41.8% to 99.7%) (**Figure S5I**). We note that some cell types identified by the original authors were poorly separated by all methods in our hand, such as Fibro-4 and Fibro-5, or “Chondro” and “Chondro-

Hyper", reflecting that the original separation was for cell subtypes or states. Furthermore, pericytes, which appeared quite distinct in our UMAP visualization and formed a distinct cluster when clustering the non-binarized data, was incorrectly mixed with fibroblasts and Osteo Lineage Cell (OLC)-2 in the clustering of binarization alone. However, this was classified correctly as a distinct cluster after applying TF/IDF and LSI to the binarized data. Taken together, these data strongly support the conclusion that binarization of the scRNA-seq data can be used to identify cell types and TF-IDF/LSI transformation improves clustering.

**Figure S5. Comparison of clustering results for a bone atlas scRNA-seq data.**
**A)**. UMAP of the non-binarized data, colored by our clusters and updated cell type annotation. The cell cluster annotation was obtained by taking the dominant cell type assignment from the original authors (see C). **B)**. Bubble plot showing the expression of the top cell type markers across clusters in A. The top two markers were selected and ordered by the marked cell types. **C)**. Heatmap matrix showing the comparison between the original authors' cell type assignment (*rows*) and our re-clusters of the non-binarized data (*columns*). **D)**. UMAP of the binarized data, colored by Louvain clustering result. **E)**. Bubble plot showing the expression of cell type markers across binarized clusters in D. **F)**. Heatmap matrix showing the comparison between the clusters of non-binarized data (*rows*) and clusters in D (*columns*). **G)**. UMAP of the binarized and TF/IDF-LSI transformed data, colored by clustering result. **H)**. Bubble plot showing the expression of cell type markers across clusters in G. **I)**. Heatmap matrix showing the comparison of clusters of non-binarized data (rows) and clusters of TF/IDF-LSI data (columns). "Lepr-MSC" = Leptin receptor positive Mesenchymal Stromal Cell; OLC = Osteo Lineage Cell; EC = Endothelial cell.

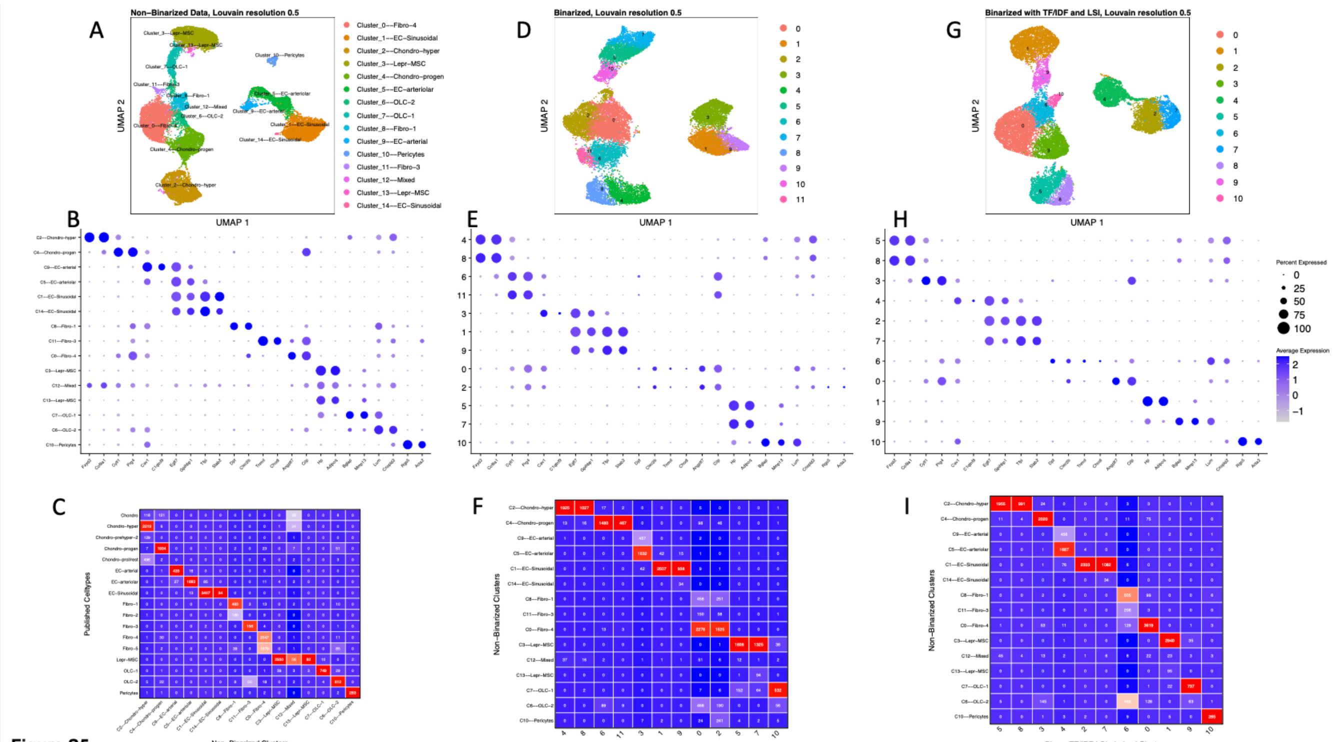


Figure S5

### 6. Cluster quality of the 10K PBMC multiomic data.

We clustered the 10K PBMC multiomic data using several approaches. QC was performed for each modality independently using the default parameters in Muon and Scanpy. Starting from an initial 11,909 cells, after QC and restricting to cells with data present for both modalities, we obtained 9,072 cells. We took the RNA portion of the data and clustered them without binarization using the standard Scanpy workflow. The first 20 PCs were used for constructing the neighborhood graph (n_neighbors=10) and Leiden clustering (resolution = 0.5). We took the ATAC portion of the data and clustered them using the workflow with default parameters for Muon as described in their documentation [7]. The first LSI component was dropped and the 2:31 components used for neighborhood graph construction (n_neighbors=10) and Leiden clustering (resolution = 0.5). We binarized the scRNA data, concatenated it with the scATAC data, performed TF-IDF transformation, LSI dimension reduction, and Leiden clustering. The first LSI component was dropped and the 2:31 components were used to calculate the neighborhood graph and Leiden clustering (resolution=1.0). We also applied MOFA to integrate the pre-binarized scRNA-seq data and scATAC-seq and then applied Leiden clustering on the integrated data. The MOFA matrix was computed using the previously assigned Highly Variable

Features in each modality. The MOFA representation was used to calculate the neighborhood graph and Leiden clustering (resolution=0.9). We obtained the cell type markers (genes or ATAC-seq peaks) from the MUON websites and examined their expression patterns in clusters obtained from each of these workflows. For bubble plots of the marker genes, the expression was based on the pre-binarized data.

**Figure S6. Expression of the PMBC cell type markers.**

**A).** Bubble plot showing the expression of the cell type markers across clusters from pre-binarized scRNA-seq portion of the multiomic data. **B).** Bubble plot showing the chromatin accessibility score of the cell type markers across clusters from scATAC-seq portion of the multiomic data. **C).** Bubble plot showing the expression of the cell type markers across clusters from concatenated pre-binarized scRNA-seq data and scATAC-seq data. **D).** Bubble plot showing the expression of the cell type markers across clusters from MOFA integration.

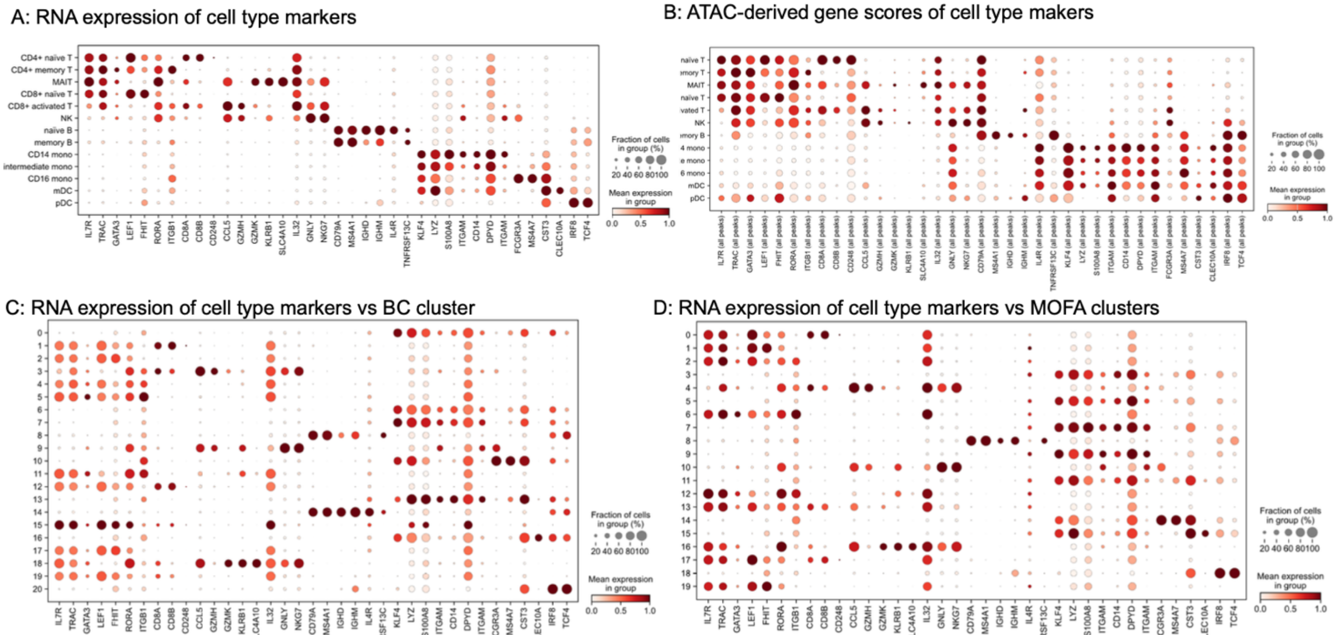

**Figure S7. Comparison of PMBC clusters from different methods.**

**A)** Clustering of the binarized scRNA-seq data, colored by clusters (left) or cell types (right)**.** **B,C).** Alluvia plot (**B**) and matrix plot (**C**) showing the cell relationship between clusters from pre-binarized scRNA-seq data and the clusters from scATAC-seq data. **D**). Matrix plot showing the cell relationship between clusters from pre-binarized scRNA-seq data and BC clusters of the integrated data. **E**). Matrix plot showing the cell relationship between clusters from scATAC-seq data and BC clusters of the integrated data. **F**). Matrix plot showing BC clusters of the integrated data and clusters of the MOFA integrated data.

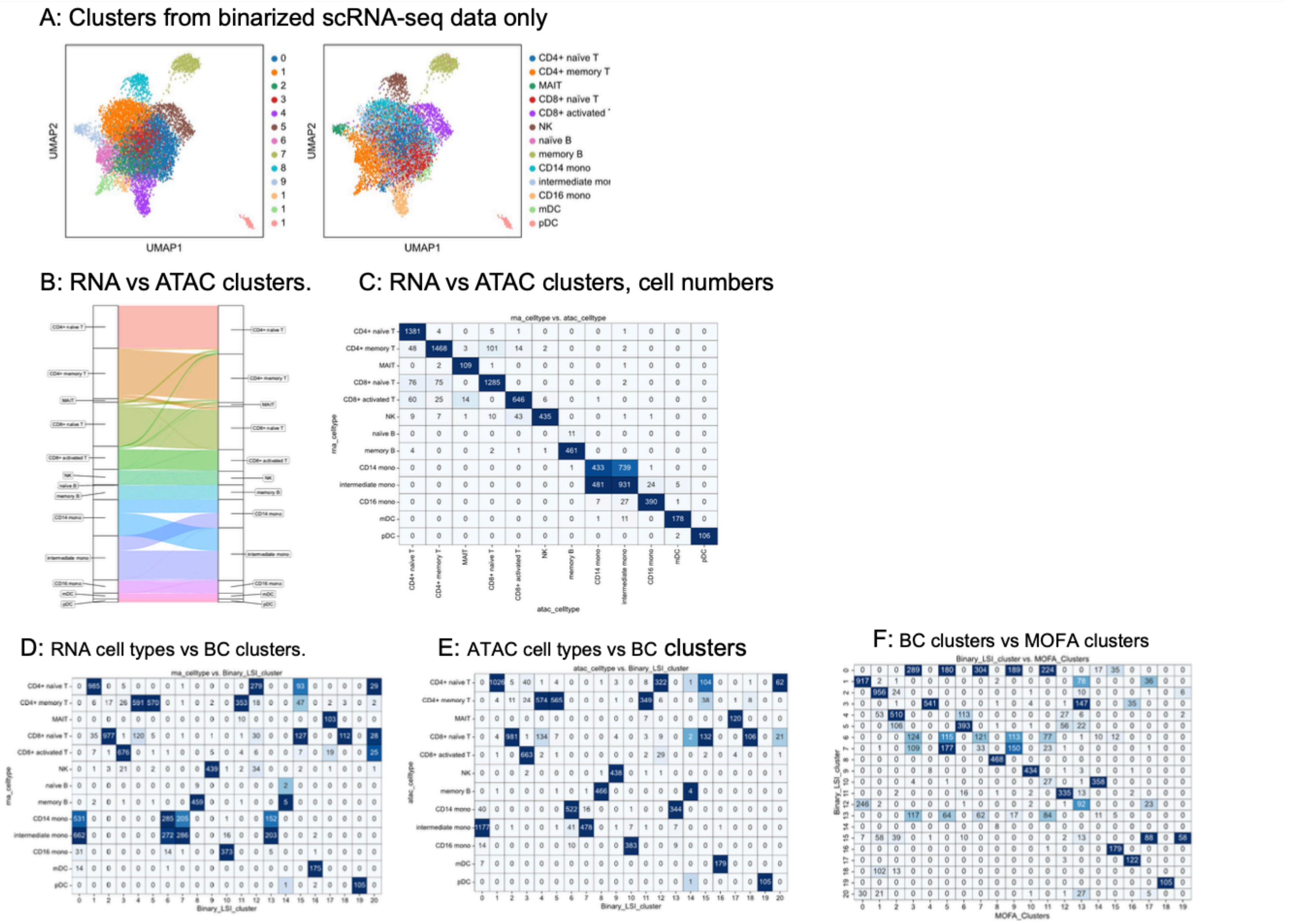


**7. Contribution of RNA and ATAC features to clustering mouse breast multiomics data.**

We obtained a multiomic data (10X platform) from a previous study that investigated cellular heterogeneity in normal breast tissue and breast cancer [8]. The study contained multiple samples from both human and mice. We analyzed 10,018 cells from only one of the samples from the normal mouse breast tissue (GSM6543819). After QC performed independently for each modality and restricting to those cells for which both scRNA-seq and scATAC-seq data were present, we obtained 9,057 cells with 25,767 genes from scRNA-seq and 93,484 fragments/peaks from scATAC-seq. Although the authors have identified the 5 major cell types, this annotation information was not available for individual cells or samples. We thus run MOFA integration using previously assigned Highly Variable Genes and Peaks, and clustering of the data (Leiden resolution = 0.1), obtaining 6 clusters, closely matching to the five cell types except T cells were split into two clusters (**Figure S8A**). We then binarized the scRNA-seq data, concatenated them with the scATAC-seq data, and clustered the cells using the top 20,000 HVFs (7,681 RNA features and 12,319 ATAC-seq features). We obtained 9 clusters. Compared to the clusters from MOFA integration, BC clustering split both endothelial and myeloid cells to 3 clusters each (**Figure S8B**). Using the MOFA integration clusters (i.e., cell types) as the reference, we tested how the ratios of RNA to ATAC features affected BC cluster accuracy (**Figure S8C,D**). As shown in **Figure S8B**, our integration seems to split two cell types into 3 clusters, which may explain the relatively low ARI scores in our comparison. To explore this, we decided to use the pre-binarized scRNA-seq data to identify markers for our 9 BC clusters. The results supported our split of the endothelial cells and suggested that the cluster 4 was likely epithelial cells that should exist in breast tissues (**Figure S8E**). Therefore, we used our 9 BC clusters from all RNA and ATAC features as the reference and re-evaluated the clustering performances from varying RNA or ATAC features. As expected, the overall performance as measured by ARI improved compared to the results using MOFA clusters as the benchmark reference (**Figure S8F,G**). While the cluster accuracies did not seem to be affected by increased RNA features (**Figure S8F**), they showed an increased and then decreased trends when more ATAC features were included (**Figure S8G**).

**Figure S8. Clustering mouse mammary parenchyma multiomics data.**
**A**). UMAPs showing the MOFA integrated data, colored by clusters (left) or cell types (right) identified using markers computed from the pre-binarized scRNA-seq data. **B**). Comparison of clusters from MOFA integration and our integration of the binarized and concatenated data, shown as Alluvia plot (left) or matrix plot (right). **C,D**). Matrix plots for the clustering accuracies when different number of RNA features (C) or ATAC features (D) were included for clustering, using the MOFA clusters as the reference (the two T cell clusters were merged for computing all these metrics). **E**). Expression of top markers for the BC clusters (C0-C8) with the corresponding cell types. The markers were computed from pre-binarized scRNA-seq data. The two UMAPs in the bottom show specific expression of *Epcam* and *Adgre1* in the C4:Endothelial-

1 and C3:Myeloid-1 cluster, indicating they are epithelia and macrophage, respectively. **F,G**). Violin plots (top) and tables (top) for the clustering accuracies when different number of RNA features (F) or ATAC features (G) were included for clustering, using our clusters from the binarized and concatenated data as the reference.

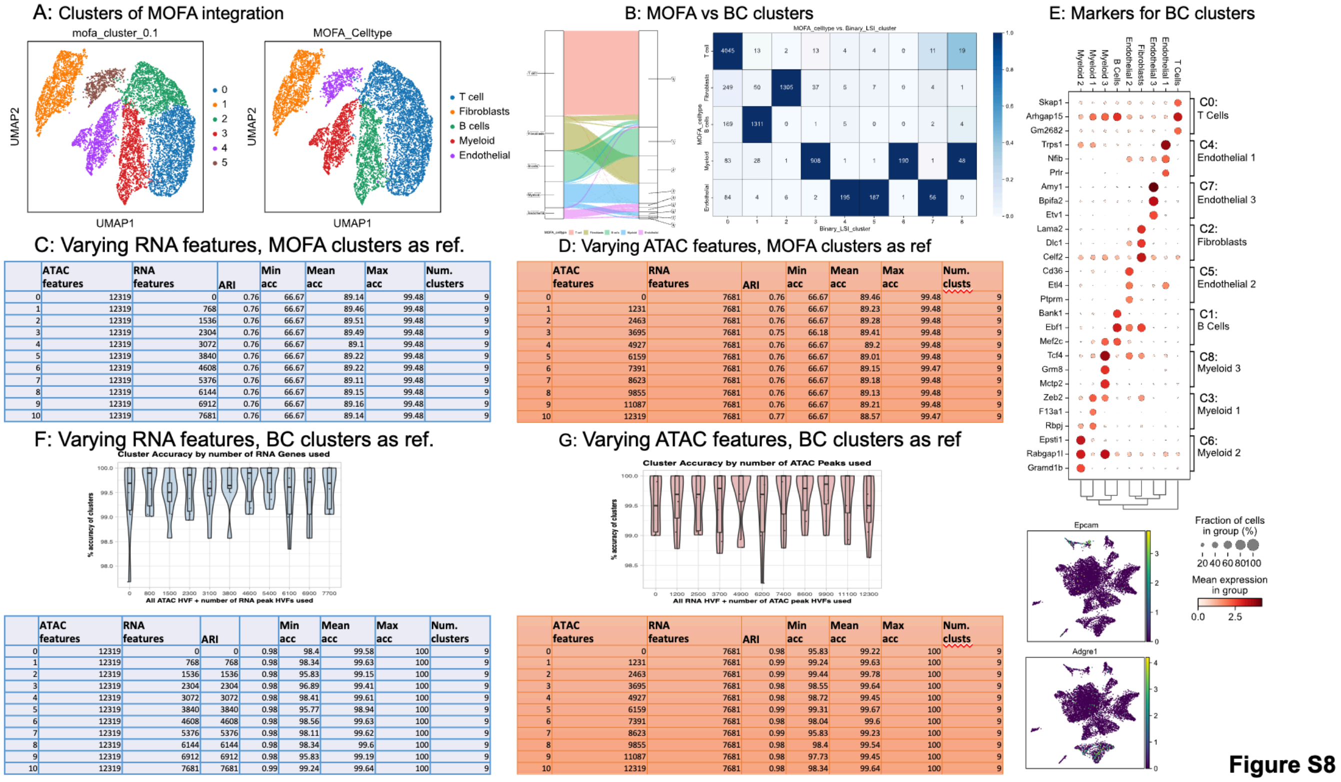


C: Varying RNA features, MOFA clusters as ref.

| | ATAC features | RNA features | ARI | Min acc | Mean acc | Max acc | Num. clusters |
|---|---|---|---|---|---|---|---|
| 0 | 12319 | 0 | 0.76 | 66.67 | 89.14 | 99.48 | 9 |
| 1 | 12319 | 768 | 0.76 | 66.67 | 89.46 | 99.48 | 9 |
| 2 | 12319 | 1536 | 0.76 | 66.67 | 89.51 | 99.48 | 9 |
| 3 | 12319 | 2304 | 0.76 | 66.67 | 89.49 | 99.48 | 9 |
| 4 | 12319 | 3072 | 0.76 | 66.67 | 89.1 | 99.48 | 9 |
| 5 | 12319 | 3840 | 0.76 | 66.67 | 89.22 | 99.48 | 9 |
| 6 | 12319 | 4608 | 0.76 | 66.67 | 89.22 | 99.48 | 9 |
| 7 | 12319 | 5376 | 0.76 | 66.67 | 89.11 | 99.48 | 9 |
| 8 | 12319 | 6144 | 0.76 | 66.67 | 89.15 | 99.48 | 9 |
| 9 | 12319 | 6912 | 0.76 | 66.67 | 89.16 | 99.48 | 9 |
| 10 | 12319 | 7681 | 0.76 | 66.67 | 89.14 | 99.48 | 9 |

D: Varying ATAC features, MOFA clusters as ref

| | ATAC features | RNA features | ARI | Min acc | Mean acc | Max acc | Num. clusts |
|---|---|---|---|---|---|---|---|
| 0 | 0 | 7681 | 0.76 | 66.67 | 89.46 | 99.48 | 9 |
| 1 | 1231 | 7681 | 0.76 | 66.67 | 89.23 | 99.48 | 9 |
| 2 | 2463 | 7681 | 0.76 | 66.67 | 89.28 | 99.48 | 9 |
| 3 | 3695 | 7681 | 0.75 | 66.18 | 89.41 | 99.48 | 9 |
| 4 | 4927 | 7681 | 0.76 | 66.67 | 89.2 | 99.48 | 9 |
| 5 | 6159 | 7681 | 0.76 | 66.67 | 89.01 | 99.48 | 9 |
| 6 | 7391 | 7681 | 0.76 | 66.67 | 89.15 | 99.47 | 9 |
| 7 | 8623 | 7681 | 0.76 | 66.67 | 89.18 | 99.48 | 9 |
| 8 | 9855 | 7681 | 0.76 | 66.67 | 89.13 | 99.48 | 9 |
| 9 | 11087 | 7681 | 0.76 | 66.67 | 89.21 | 99.48 | 9 |
| 10 | 12319 | 7681 | 0.77 | 66.67 | 88.57 | 99.47 | 9 |

F: Varying RNA features, BC clusters as ref.

| | ATAC features | RNA features | ARI | | Min acc | Mean acc | Max acc | Num. clusters |
|---|---|---|---|---|---|---|---|---|
| 0 | 12319 | 0 | 0 | 0.98 | 98.4 | 99.58 | 100 | 9 |
| 1 | 12319 | 768 | 768 | 0.98 | 98.34 | 99.63 | 100 | 9 |
| 2 | 12319 | 1536 | 1536 | 0.98 | 95.83 | 99.15 | 100 | 9 |
| 3 | 12319 | 2304 | 2304 | 0.98 | 96.89 | 99.41 | 100 | 9 |
| 4 | 12319 | 3072 | 3072 | 0.98 | 98.41 | 99.61 | 100 | 9 |
| 5 | 12319 | 3840 | 3840 | 0.98 | 95.77 | 98.94 | 100 | 9 |
| 6 | 12319 | 4608 | 4608 | 0.98 | 98.56 | 99.63 | 100 | 9 |
| 7 | 12319 | 5376 | 5376 | 0.98 | 98.11 | 99.62 | 100 | 9 |
| 8 | 12319 | 6144 | 6144 | 0.98 | 98.34 | 99.6 | 100 | 9 |
| 9 | 12319 | 6912 | 6912 | 0.98 | 95.83 | 99.19 | 100 | 9 |
| 10 | 12319 | 7681 | 7681 | 0.99 | 99.24 | 99.64 | 100 | 9 |

G: Varying ATAC features, BC clusters as ref

| | ATAC features | RNA features | ARI | Min acc | Mean acc | Max acc | Num. clusts |
|---|---|---|---|---|---|---|---|
| 0 | 0 | 7681 | 0.98 | 95.83 | 99.22 | 100 | 9 |
| 1 | 1231 | 7681 | 0.99 | 99.24 | 99.63 | 100 | 9 |
| 2 | 2463 | 7681 | 0.99 | 99.44 | 99.78 | 100 | 9 |
| 3 | 3695 | 7681 | 0.98 | 98.55 | 99.64 | 100 | 9 |
| 4 | 4927 | 7681 | 0.98 | 98.72 | 99.45 | 100 | 9 |
| 5 | 6159 | 7681 | 0.99 | 99.31 | 99.67 | 100 | 9 |
| 6 | 7391 | 7681 | 0.98 | 98.04 | 99.6 | 100 | 9 |
| 7 | 8623 | 7681 | 0.99 | 95.83 | 99.23 | 100 | 9 |
| 8 | 9855 | 7681 | 0.98 | 98.4 | 99.54 | 100 | 9 |
| 9 | 11087 | 7681 | 0.98 | 97.73 | 99.45 | 100 | 9 |
| 10 | 12319 | 7681 | 0.98 | 98.34 | 99.64 | 100 | 9 |

**Figure S9. UMAPs showing improved separation of specific breast cell types with increasing RNA or ATAC features for BC clustering.**

**A-D**), UMAPs showing clusters from increasing RNA features. With increasing numbers of added RNA features, the endothelial cells became more separated, with the potential epithelial cluster (#4, endothelial-1, *Epcam+*/*Pecam1*-) being further separated from other endothelial cells. These clusters were maximally separated in C. Similarly, myeloid cells became more distinct with increasing RNA features. **E-H**), UMAPs showing clusters from increasing ATAC features. In RNA only (E), myeloid were already well separated into at least two distinct clusters (3 and 6), where cluster 3 was likely macrophages expressing high *Adgre1* and *Csf1r*. This separation was reduced with increasing ATAC features.

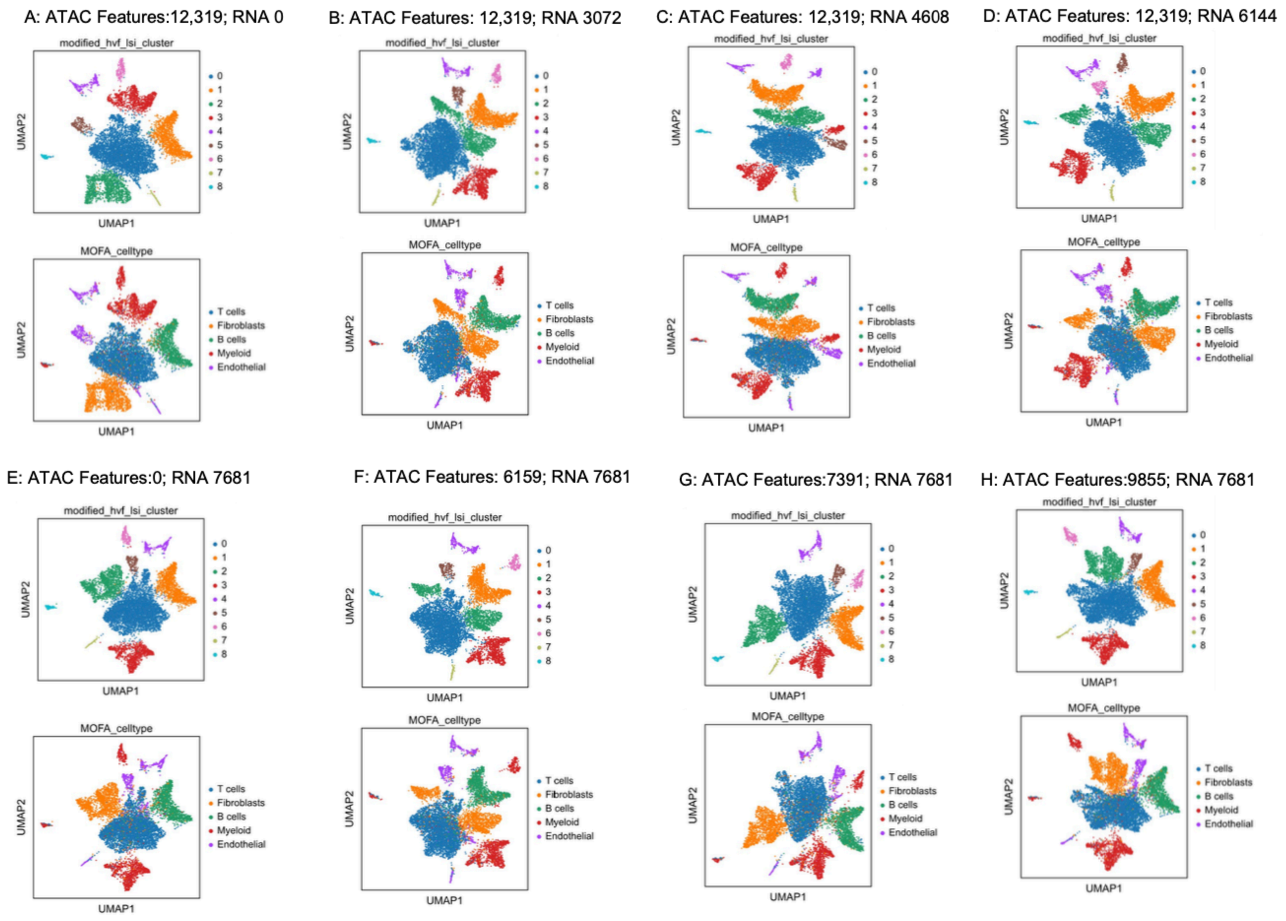


**8. Contribution of RNA and ATAC features to clustering human brain cortex multiomics data.**

To study how the numbers of RNA or ATAC features could affect the clustering results of the human brain cortex multiomics data, we binarized the scRNA-seq data, concatenated them with the scATAC-seq data, and then used Seurat and Signac to carry out TF/IDF and LSI. After inspection of elbow plots, the LSI components 2 to 20 were used for Louvain clustering with resolution set to 0.2. The cluster results were compared to the authors' cell type annotation (**Figure S10A**). We found surprisingly that one (#10) of our clusters contained cells from all cell types, while for the other clusters the majority of the cells were from a single cell type. To investigate this, we analyzed the expression pattern of the original authors' cell type markers in either all cells excluding cluster 10 cells (**Figure S10B**) or only cells in our cluster 10 (**Figure S10C**). The results showed that the cells in cluster 10 did not express the cell type markers that were expressed in the correct cell types as described previously [9]. We further analyzed the number of total RNA transcripts (nCount_RNA), the number of unique genes detected (nFeature_RNA), and the %s of mitochondrial UMIs in our integrated clusters (**Figure S10D**). The data indicated that the cluster 10 was composed of poor-quality cells or droplets with low numbers of UMIs and unique genes. This explains why such distinct cell types were classified together in cluster 10 after data binarization. To further support this, we computed cluster markers using the pre-binarized scRNA-seq data and found that cluster 10 had hardly any specific markers (**Figure S10E**). This result suggests that binarization may help to identify and remove noise cells in some cases.

Next, we performed an analysis to compare the inclusion ratio of two feature types. First, we used all ATAC features, while incrementally increasing the number of binarized RNA features sorted by their variances, and then performed TF/IDF-LSI and clustering. We kept other analytic parameters the same across increments, including number of LSI components (2:20) and Louvain clustering resolution (0.2). After that, we determined the cluster accuracies using the authors' cell type annotation as reference. The results (**Figure S10F**) indicate that adding ATAC features could increase clustering performance. Similarly, we fixed the number of RNA features (n = 2000 top HVFs), gradually included the numbers of ATAC features sorted by the numbers of reads in all cells, and then performed clustering. The results (**Figure S10G**) indicate that adding RNA features did not systematically affect clustering performance but affected some clusters more than others. More specifically, endothelial cells were closely clustered with vascular smooth muscle cells (VSMC) and pericytes with no RNA features, but they became to separate from VSMC and pericytes as more RNA features were added. VSMC and pericytes, however, were still co-clustered. When RNA features were fixed, the oligodendrocyte progenitors (OPC) cluster were split into two well separated clusters with more ATAC features added; the three inhibitory neurons (IN-MGE / IN-CGE / IN-Fetal) also became to form more distinct clusters with more ATAC features.

**Figure S10. Clustering human brain cortex multiomics data.**

**A**). Comparison of published cell types and our integrated clusters from full data. **B**). Expression of cell type markers in all cells excluding the cluster 10 cells. **C**). Expression of cell type markers in cells in the cluster 10 only. **D**). Quality metrics of cells based on RNA data. **E**). Expression of markers computed for the clusters in our integration. **F**). Clustering performance using fixed number of ATAC features but varying RNA features. **G**). Clustering performance using fixed number of RNA features but varying ATAC features.

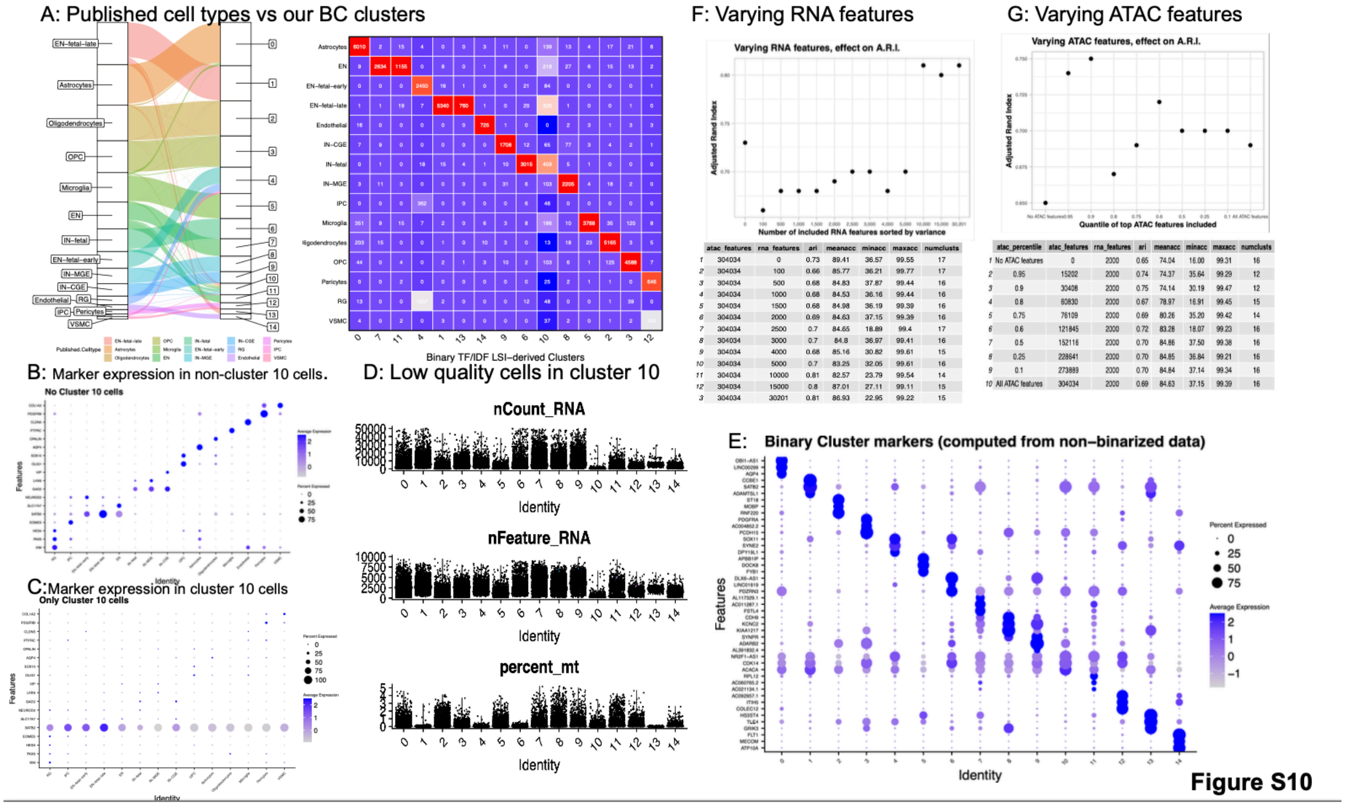


| | atac_features | rna_features | ari | meanacc | minacc | maxacc | numclusts |
|---|---|---|---|---|---|---|---|
| 1 | 304034 | 0 | 0.73 | 89.41 | 36.57 | 99.55 | 17 |
| 2 | 304034 | 100 | 0.66 | 85.77 | 36.21 | 99.77 | 17 |
| 3 | 304034 | 500 | 0.68 | 84.83 | 37.87 | 99.44 | 16 |
| 4 | 304034 | 1000 | 0.68 | 84.53 | 36.16 | 99.44 | 16 |
| 5 | 304034 | 1500 | 0.68 | 84.98 | 36.19 | 99.39 | 16 |
| 6 | 304034 | 2000 | 0.69 | 84.63 | 37.15 | 99.39 | 16 |
| 7 | 304034 | 2500 | 0.7 | 84.65 | 18.89 | 99.4 | 17 |
| 8 | 304034 | 3000 | 0.7 | 84.8 | 36.97 | 99.41 | 16 |
| 9 | 304034 | 4000 | 0.68 | 85.16 | 30.82 | 99.61 | 15 |
| 10 | 304034 | 5000 | 0.7 | 83.25 | 32.05 | 99.61 | 16 |
| 11 | 304034 | 10000 | 0.81 | 82.57 | 23.79 | 99.54 | 14 |
| 12 | 304034 | 15000 | 0.8 | 87.01 | 27.11 | 99.11 | 15 |
| 3 | 304034 | 30201 | 0.81 | 86.93 | 22.95 | 99.22 | 15 |

| | atac_percentile | atac_features | rna_features | ari | meanacc | minacc | maxacc | numclusts |
|---|---|---|---|---|---|---|---|---|
| 1 | No ATAC features | 0 | 2000 | 0.65 | 74.04 | 16.00 | 99.31 | 16 |
| 2 | 0.95 | 15202 | 2000 | 0.74 | 74.37 | 35.64 | 99.29 | 12 |
| 3 | 0.9 | 30408 | 2000 | 0.75 | 74.14 | 30.19 | 99.47 | 12 |
| 4 | 0.8 | 60830 | 2000 | 0.67 | 78.97 | 16.91 | 99.45 | 15 |
| 5 | 0.75 | 76109 | 2000 | 0.69 | 80.26 | 35.20 | 99.42 | 14 |
| 6 | 0.6 | 121845 | 2000 | 0.72 | 83.28 | 18.07 | 99.23 | 16 |
| 7 | 0.5 | 152116 | 2000 | 0.70 | 84.86 | 37.50 | 99.38 | 16 |
| 8 | 0.25 | 228641 | 2000 | 0.70 | 84.85 | 36.84 | 99.21 | 16 |
| 9 | 0.1 | 273889 | 2000 | 0.70 | 84.84 | 37.14 | 99.34 | 16 |
| 10 | All ATAC features | 304034 | 2000 | 0.69 | 84.63 | 37.15 | 99.39 | 16 |

**Figure S11. UMAPs showing improved separation of some brain cell types with increasing RNA or ATAC features.**

**A-C**), UMAPs showing clusters from increasing RNA features. Green arrows indicate position of endothelial cell clusters. **D-G**), UMAPs showing clusters from increasing ATAC features. Lime green arrows indicate positions of IN-MGE, IN-CGE, and IN-Fetal clusters. Purple arrows indicate position of OPC cluster(s).

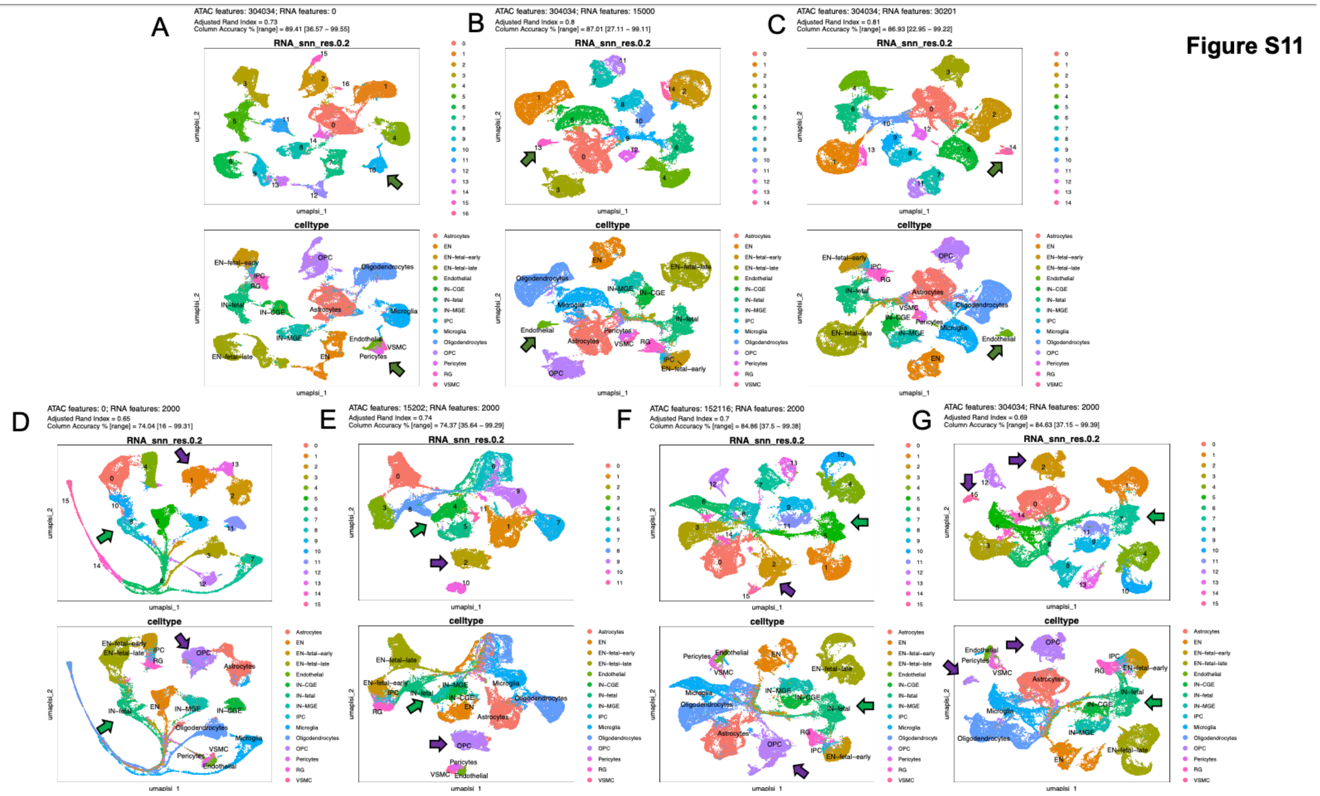

**References**


1. Processing gene expression of 10k PBMCs — muon-tutorials documentation. https://muon-tutorials.readthedocs.io/en/latest/single-cell-rna-atac/pbmc10k/1-Gene-Expression-Processing.html.
2. Polański, K. *et al.* BBKNN: fast batch alignment of single cell transcriptomes. *Bioinformatics* **36**, 964–965 (2020).
3. Baryawno, N. *et al.* A Cellular Taxonomy of the Bone Marrow Stroma in Homeostasis and Leukemia. *Cell* **177**, 1915-1932.e16 (2019).
4. Tarhan, L. *et al.* Single Cell Portal: an interactive home for single-cell genomics data. 2023.07.13.548886 Preprint at https://doi.org/10.1101/2023.07.13.548886 (2023).
5. Butler, A., Hoffman, P., Smibert, P., Papalexi, E. & Satija, R. Integrating single-cell transcriptomic data across different conditions, technologies, and species. *Nat. Biotechnol.* **36**, 411–420 (2018).
6. Hafemeister, C. & Satija, R. Normalization and variance stabilization of single-cell RNA-seq data using regularized negative binomial regression. *Genome Biol.* **20**, 296 (2019).
7. Processing chromatin accessibility of 10k PBMCs — muon-tutorials documentation. https://muon-tutorials.readthedocs.io/en/latest/single-cell-rna-atac/pbmc10k/2-Chromatin-Accessibility-Processing.html.
8. Foster, D. S. *et al.* Multiomic analysis reveals conservation of cancer-associated fibroblast phenotypes across species and tissue of origin. *Cancer Cell* **40**, 1392-1406.e7 (2022).
9. Zhu, K. *et al.* Multi-omic profiling of the developing human cerebral cortex at the single-cell level. *Sci. Adv.* **9**, eadg3754.
